# A novel combination of Class-I fumarases and metabolites (α-ketoglutarate and fumarate) signal the DNA damage response in *E. coli*

**DOI:** 10.1101/2020.08.04.232652

**Authors:** Yardena Silas, Esti Singer, Koyeli Das, Norbert Lehming, Ophry Pines

**Affiliations:** Department of Microbiology and Molecular Genetics, IMRIC, Faculty of Medicine, Hebrew University, Jerusalem, Israel; CREATE⍰NUS⍰HUJ Program and the Department of Microbiology and Immunology, Yong Loo Lin School of Medicine, National University of Singapore, Singapore, Singapore

**Keywords:** Fumarase, α-KG, DNA damage, *Escherichia coli*, AlkB

## Abstract

Class-II fumarases (Fumarate Hydratase, FH) are dual targeted enzymes, occurring in the mitochondria and cytosol of all eukaryotes. They are essential components in the DNA damage response (DDR) and more specifically, protecting cells from DNA double strand breaks. Similarly, the Gram-positive Bacterium *Bacillus subtilis* Class-II fumarase, in addition to its role in the TCA cycle, also participates in the DDR. *Escherichia coli*, harbors three fumarase genes; Class-I *fumA* and *fumB* and Class-II *fumC*. Notably, Class-I fumarases, show no sequence similarity to Class-II fumarases and are of different evolutionary origin. Strikingly, here we show that *E. coli* fumarase functions are distributed between Class-I fumarases which participate in the DDR, and the Class-II fumarase which participates in respiration. In *E. coli*, we discover that the signaling molecule, alpha-ketoglutarate (α-KG), has a novel function, complementing DNA damage sensitivity of *fum* null mutants. Excitingly, we identify the *E. coli* α-KG dependent DNA repair enzyme AlkB, as the target of this interplay of metabolite signaling. In addition to α-KG, fumarate (fumaric acid) is shown to affect DNA damage repair on two different levels, first by directly inhibiting the DNA damage repair enzyme AlkB demethylase activity, both in vitro and in vivo (countering α-KG). The second is a more global effect on transcription, as *fum* null mutants exhibit a decrease in transcription of key DNA damage repair genes. Together these results show evolutionary adaptable metabolic signaling of the DDR, in which fumarases and different metabolites are recruited regardless of the evolutionary enzyme Class preforming the function.

**Significance Statement:** Class-II fumarases have been shown to participate in cellular respiration and the DNA damage response. Here we show, for the first time, that in the model prokaryote, *Escherichia coli*, which harbors both Class-I and Class-II fumarases, it is the Class-I fumarases that participate in DNA damage repair by a mechanism which is different than those described for other fumarases. Strikingly, this mechanism employs a novel signaling molecule, alpha-ketoglutarate (α-KG), and its target is the DNA damage repair enzyme AlkB. In addition, we show that fumarase precursor metabolites, fumarate and succinate, can inhibit the α-KG-dependent DNA damage repair enzyme, AlkB, both in vitro and in vivo. This study provides a new perspective on the function and evolution of metabolic signaling.

## Introduction

Fumarase (fumarate hydratase, FH) is an enzyme that participates in the tricarboxylic acid (TCA) cycle, there, it catalyzes the reversible hydration of fumarate to L-malate [1]. In eukaryotes, in addition to its mitochondrial localization, a common theme conserved from yeast to humans is the existence of a cytosolic form of fumarase [2, 3]. The dual distribution of fumarase between the mitochondria and the cytosol is a universal trait, but intriguingly, the mechanism by which dual distribution occurs does not appear to be conserved [4–8]. We discovered that in eukaryotes the cytosolic form of this TCA enzyme was shown to have an unexpected function in the DNA damage response (DDR), and specifically, a role in recovery from DNA double strand breaks (DSBs) [9]. It appears that metabolic signaling in these organisms is achieved via the fumarase organic acid substrate, fumarate [10, 11]. A recent study by Singer et al. [12] shows that the Gram-positive *Bacillus subtilis* fumarase, has a role in the TCA cycle, as well as a role in the DDR. Fumarase dependent signaling of the DDR in that bacterium is achieved by its metabolic product, L-malate, which affects RecN (the first enzyme recruited to DNA damage sites) at the level of expression and localization. That study indicated, that the dual function of this enzyme predated its dual distribution and, according to our model, was the driving force for the evolution of dual targeting in eukaryotes [12].

An intriguing variation to the themes above, is the fact that there are two distinct classes of fumarase. In the organisms mentioned above (*Saccharomyces cerevisiae*, human, *B. subtilis*) the fumarase that participates in both the TCA cycle and in DNA repair, belongs to the Class-II fumarases. In contrast, the Class-I fumarases, which bear no sequence or structural similarity to Class-II fumarases, are predominantly found in prokaryotes [1] and in early evolutionary divergents of the eukaryotic kingdom (protozoa and invertebrates). The model organism that we have employed in this study is the Gram-negative bacterium *E. coli* which harbors three fumarase genes; Class-I heat-labile, iron-sulfur cluster containing fumarases, *fumA* and *fumB*, and Class-II heat-stable, *fumC*. FumA and FumB are homologous proteins, sharing 90% amino acid sequence identity. Both FumA and FumB kinetic parameters are very similar with respect to the natural substrates L-malate and fumarate [13]. FumC is homologous to eukaryotic and *B. subtilis* fumarases, sharing 64% amino acid sequence identity. *E. coli* fumarase expression is aerobically controlled; FumB expression was found to be four-fold elevated under anaerobic conditions [14], with FumA peak expression during normal aerobic growth. FumC was found to express weakly under aerobic conditions, and appears to be the backup enzyme, synthesized mainly when iron is low [15].

Employing the *E. coli* model, we ask whether Class-I and/or Class-II fumarases are involved in the DDR and/or the TCA cycle and what is the distribution of functions between these two Classes of fumarases in this organism. Here we show that FumA and FumB are participants of the DDR in *E. coli*, while FumC naturally participates in cellular respiration. FumA and FumB participation in the DDR is based on a fascinating interplay of TCA cycle Intermediates; alpha-ketoglutarate (α-KG), fumarate and succinate affecting the activity of the α-KG dependent DNA repair enzyme, AlkB, which is required for successful repair of methylated/alkylated damaged DNA. We also show, that in *E. coli*, the absence of fumarase affects the transcription of many genes following DNA damage induction, including DNA repair genes.

## Results

### *ΔfumA* and *ΔfumB* but not *ΔfumC E. coli* strains, are sensitive to DNA damage induction

In order to examine a possible role for *E. coli* fumarases in the DNA damage response, we induced DNA damage in *E. coli* strains lacking each of the fumarase genes, by exposing them to treatment with ionizing radiation (IR, Fig. 1A) or methyl methanesulfonate (Fig. 1B, MMS, 0.35% [v/v] for 30 or 45 minutes). Figures 1A and 1B (compare rows 2 and 3 in each panel to row 1) demonstrate that compared to the control (WT), strains lacking *fumA* or *fumB* are very sensitive to DNA damage (200 Gy [IR] or 0.35% [v/v] MMS for 45 minutes), while the strain lacking *fumC* appears to be unaffected by the DNA damage induction (row 4). These results suggest that FumA and FumB play a role in the DNA damage response, while FumC has no observable role in this process. While the levels of FumA exhibit a very slight insignificant change in protein levels upon treatment with MMS (Fig 1C top panel), the levels of FumC increase (Fig 1C second panel), thereby revealing a more complex picture of regulation. Intriguingly, the double *fumA/fumB* mutant (Fig 1D, middle panel, *ΔfumAB*, Row 5) is resistant to DNA damage induction, either IR (SI Appendix, Fig. S3B) or MMS treatment and shows a phenotype resembling that of the WT strain (Row 1). Compare to single *fumA* and *fumB* deletion strains (*ΔfumA, ΔfumB*, rows 2 and 3) or a triple mutant lacking all three fumarase genes (*ΔfumACB*, row 6) which are sensitive to IR (SI Appendix, Fig. S3B) and MMS. One possible explanation for this phenomenon is that in the absence of FumA and FumB, FumC expression in the *ΔfumAB* strain (i.e., no MMS treatment) increases to undertake both roles of respiration and participation in the DDR (Fig 1E). Worth mentioning here and referred to in the next section, is that complementation of the DNA damage sensitivity can be achieved by expression of the genes in trans (SI Appendix, Fig. S1A,B).

**Figure 1.**
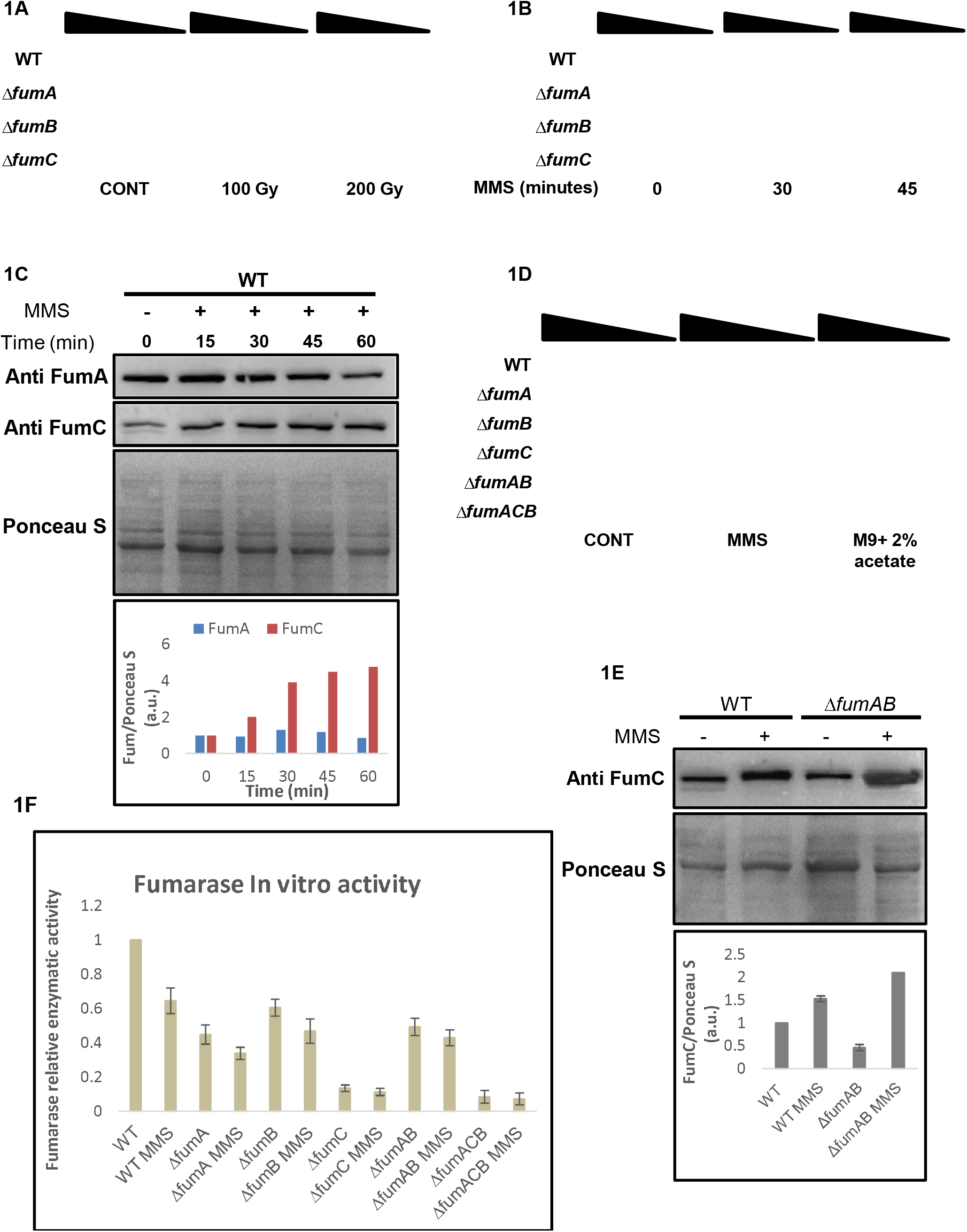
*E. coli* FumA and FumB are required for DNA damage repair. (**A, B**) *E. coli* wild type (WT), *ΔfumA, ΔfumB, ΔfumC*, strains were grown to mid-exponential phase (OD600nm=0.3), irradiated (100 / 200 Gy) <u>or</u> treated with MMS (0.35% (v/v) for 30 or 45 minutes at 37□C) respectively. The cells were then washed and serially diluted (spot test) onto LB plates. (**C**) *E. coli* wild type (WT) was treated with MMS (0.35% [v/v] for 45 minutes at 37□C), the cells were lysed and centrifuged to obtain a supernatant which was subjected to Western blotting (using the indicated antibodies) and Ponceau S staining. The chart presents the relative amount of FumA and FumC, with or without MMS according to densitometric analysis of Fig. 1C and normalized to Ponceau S staining (**D**) *E. coli* strains were grown and subjected to spot assay as in Fig. 1B (n=3, for all spot test assays) (**E**) *E. coli* WT and *ΔfumAB* were grown as in 1B (0.35% MMS (v/v) for 45 min) and subjected to protein extraction and Western blotting. The chart presents the relative amount of FumC, with or without MMS according to densitometric analysis of Fig. 1E and normalized to Ponceau S staining (mean ±SEM [n = 3], p<0.05). (**F**) *E. coli* strains were grown to midexponential phase, the cells were lysed and centrifuged to obtain the supernatant which was assayed for fumarase activity at 250 nm with L-malic acid as the substrate (mean ± SEM [n = 3]).

### All three *E coli* fumarase genes can participate in the TCA cycle

It is important to emphasize, with regard to respiration, that a functional TCA cycle including participation of fumarase is required (Figure 1D, right panel). Accordingly, we find growth of our strains (WT, *ΔfumA, ΔfumB, ΔfumC* and *ΔfumAB*) which harbor at least one functional fumarase gene, on minimal media with acetate as a sole carbon and energy source (requiring cellular respiration for growth). In contrast, the triple mutant (*ΔfumACB*), completely lacking fumarase, is unable to respire and therefore unable to grow on this media (Figure 1D, right panel row 6). The Δ*fumC* mutant shows an interesting respiration phenotype, although FumA and/or FumB seem to participate primarily in the DDR, it would seem that in the absence of FumC, these enzymes have the ability to participate in respiration, but likely play a less critical role in this function. Figure 1F shows the relative enzymatic activity of each of the mutant strains compared to the WT. These strains were assayed for fumarate production using L-malate as a substrate, testing its depletion at 250nm. We find an observable decrease in enzymatic activity in fumarase mutant strains when compared to the WT (*ΔfumA* 50%, *ΔfumB* 30%, *ΔfumC* 90%, *ΔfumAB* 50% and *ΔfumACB* with insignificant activity). After treatment with MMS, the WT exhibits a 40% reduction in enzymatic activity and the mutants also exhibit an additional reduction in enzymatic activity (e.g. *ΔfumA 10%, ΔfumB 15%, ΔfumAB 5%*) when compared to the same untreated strain. The sharp decrease of fumarase activity in *ΔfumC*, indicates that FumC is responsible for most of the fumarase activity in *E. coli*. FumA and FumC protein levels of these strains under the same conditions are shown in figure S1A. These strains also show an expected decrease in FumC protein levels in the mutants compared to the WT strain, coupled with further reduction in FumC levels following MMS treatment (with the exception of the Δ*fumAB* strain, as referred to above). FumA levels in Δ*fumB* and Δ*fumC* strains resemble the FumA levels observed in the WT strain before induction of DNA damage, but show more significant reduction in FumA levels compared to the WT strain following MMS treatment.

As controls for the above experiments, we show that over-expression of FumA, FumB or FumC from high copy number plasmids (pUC-18) in the triple fumarase mutant (*ΔfumACB*) brings about a partial complementation of DNA damage sensitivity while only FumC appears to fully complement respiration (SI Appendix, Fig. S1A). The expression levels of each of these constructs is shown in SI Appendix, Fig. S1B. Due to the fact that these experiments are not under natural induction, but rather under over-expression conditions, FumC seems to have the same effect as FumA and FumB in regard to DNA damage induction. Nevertheless, it is also evident that under conditions of *fumA* and *fumB* absence, FumC, which is then overexpressed, has the capacity to participate in the DDR.

### *E. coli* fumarases function in both the DDR and respiration in a yeast model system

In order to validate that indeed FumA and FumB participate in the DNA damage response and FumC participates in respiration, we employed, as in the past [9, 12], the yeast *S. cerevisiae* as a model. We cloned each of these genes into yeast expression vectors and transformed them into two different yeast strains. The first yeast strain, Fum1M, harbors a chromosomal *fum1* deletion (*Δfum1*) and a *FUM1* open reading frame (ORF) insertion in the mitochondrial DNA. This allows exclusive mitochondrial fumarase expression, lacking extra-mitochondrial (cytosolic/nuclear) fumarase [9]. This Fum1M strain, which is sensitive to DNA damage, was transformed with each of the fumarases separately (Fig. 2A), in order to test whether bacterial FumA and FumB are able to protect against DNA damage, a function of the cytosolic yeast fumarase. To this end, we induced DNA damage by hydroxyurea (HU) as shown in Fig. 2A (or by IR [200Gy], SI Appendix, Fig. S2A). Compared to the WT and Fum1M strains (rows 1 and 2 respectively), *E. coli* FumA and FumB are able to protect yeast from DNA damage while FumC does not (rows 3, 4 and 5 respectively).

**Figure 2.**
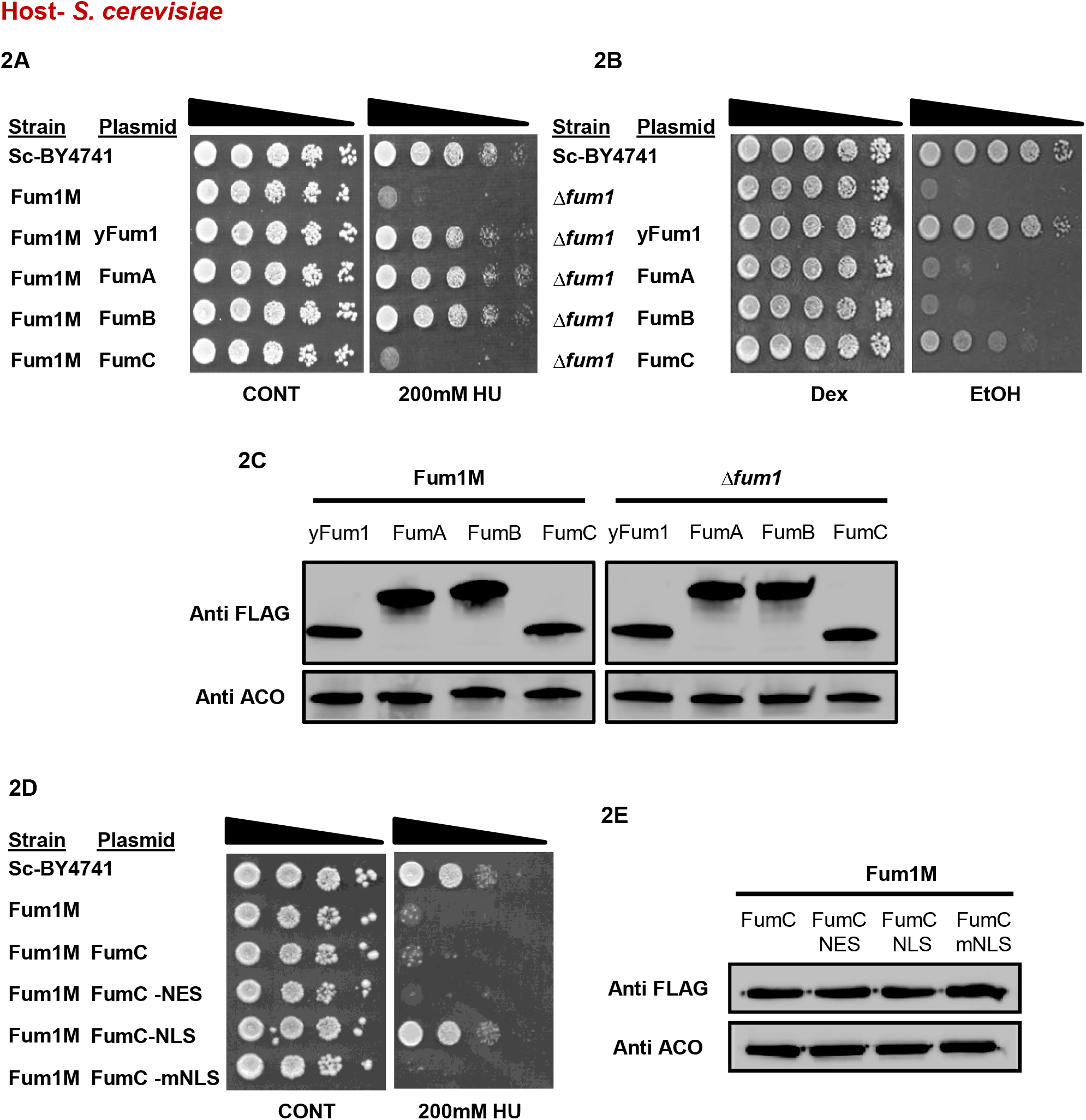
FumA, FumB and FumC substitute yeast Fum1 functions in the S. cerevisiae model system. (**A**) *S. cerevisiae* wild type BY4741 (Sc-BY4741), Fum1M and Fum1M harboring plasmids encoding the indicated *S. cerevisiae* and *E. coli fum* genes, were grown in galactose synthetic complete (SC-Gal) medium, irradiated and serially diluted onto dextrose synthetic complete (SC-Dex) plates. (**B**) *S. cerevisiae* wild type Sc-BY4741, *Δfum1* and *Δfum1* strains harboring plasmids encoding the indicated *S. cerevisiae* and *E. coli fum* genes, were serially diluted (spot test) onto dextrose or ethanol plates. (**C**) Strains harboring plasmids encoding indicated proteins (yFum1, FumA, FumB, FumC) were lysed and centrifuged to obtain the supernatant, and extracts were subjected to Western blotting, using the indicated antibodies (anti Aco1 is the loading control). (**D**) *S. cerevisiae* wild type Sc-BY4741, Fum1M and Fum1M harboring plasmids encoding the indicated plasmids, were grown to exponential phase in SC-Gal medium, serially diluted (spot test) onto SC-Dex or SC-Dex + Hydroxyurea (200 mM, HU). (**E**) Strains harboring plasmids encoding variant recombinant *fumC* genes were lysed and centrifuged to obtain the supernatant, and extracts were subjected to Western blotting, using the indicated antibodies (anti Aco1 is the loading control). Each result is representative of three independent experiments.

**Figure 3.** FumA and FumB catalytic activity is required for their DNA damage related function. (**A**) Model structure as proposed by I-TASSER, catalytic site (homologous to *L. major* active site) as proposed by PEPTIMAP, and location of important amino acids for the function of this enzyme proposed by CONSURF. (**B**) Alignment of proposed FumA/FumB structure with Fumarate Hydratase of *Leishmania major* (5l2R). (**C**) *E. coli* wild type (WT), *ΔfumA* and *ΔfumA* harboring the indicated plasmids and (**D**) *E. coli* wild type (WT), *ΔfumB* and *ΔfumB* harboring the indicated plasmids were grown to mid-exponential phase (OD600nm=0.3), treated with MMS (0.35% MMS (v/v) for 45 min) and subjected to a spot test assay. (**E**) *E. coli ΔfumA* and *ΔfumB* harboring the indicated plasmids were grown to mid-exponential phase (OD600nm=0.3) and treated with MMS (0.35% MMS (v/v) for 45 min). The cells were lysed and centrifuged to obtain the supernatants which were subjected to Western blotting using the indicated antibodies. (**F, G**) *fum* null mutant (*ΔfumACB*) harboring the indicated plasmids were grown to exponential phase, the cells were lysed and centrifuged to obtain the supernatant which was assayed for fumarase activity at 250 nm with L-malic acid as the substrate (mean ± SD [n = 3], two-tailed *t*-test *p < 0.05; **p < 0.01).

The second yeast strain completely lacks fumarase (Δ*fum1*) and therefore lacks respiration ability. We transformed each of the fumarase genes separately (Fig. 2B) and growth was followed on synthetic complete (SC) medium supplemented with ethanol (as a sole carbon and energy source, requiring cellular respiration for growth). Compared to the Δ*fum1* strain that lacks respiration ability (Fig. 2B, compare row 2 to row 1), and to the Δ*fum1* strain that harbors yeast Fum1 encoded from a plasmid (row 3), FumA and FumB are unable to complement respiration, while FumC partially complements respiration (rows 4, 5 and 6 respectively). The expression levels of each of these constructs is similar relative to the loading control aconitase (Fig. 2C). As FumA and FumB harbor no structural similarity to the yeast fumarase, this result does not definitively mean that they do not participate in respiration, but that due to evolutionary structural incompatibility with other cellular machineries, these enzymes are not able to complement respiration in this higher organism.

Previously published work from our lab [16], demonstrated that fusion of the *fumC* coding sequence to *Neurospora crassa* F_o_-ATPase, Su9 mitochondrial targeting sequence (MTS), led to complete import and processing of *E. coli* FumC in mitochondria. Without this sequence, FumC is not targeted and imported into the mitochondria, but can (complement yeast *fum* null mutants in cell respiration due to transport of the organic acids (fumarate and L-malate) in and out of this organelle. The complementation of DNA damage sensitivity by FumA and FumB indicates that these enzymes indeed participate in the DDR and this function is evolutionary conserved [12]. FumC is homologous to eukaryotic and *B. subtilis* fumarases, and as referred to above, the cytosolic role of these fumarases is in the DDR. Intriguingly, in *E. coli* this role is carried out by Class-I fumarases, although they have no sequence or structural similarity to the Class-II fumarases, besides their activity. To challenge this apparent distribution of functions between *E. coli* Class-I and Class-II fumarases and, in particular, the inability of FumC to function in the DDR, we asked whether this is due to a problem in the localization of FumC to the nucleus. Previous research shows that upon DNA damage induction there is migration of fumarase into the nucleus, there it participates in various DNA damage repair pathways [9, 10]. The approach we took was to force the FumC protein into the yeast nucleus. We constructed a *fumC* ORF fused to a nuclear localization sequence (NLS). The modified protein was expressed in a yeast Fum1M strain and the respective strains were grown with or without HU (Fig 2D). We found that the modified FumC, containing an NLS was capable of complementing the Fum1M sensitivity to HU (Fig 2D, row 5). As controls we show that the original FumC, a FumC that harbors a nuclear export signal (NES) (row 4) or a FumC harboring a mutated NLS (mNLS, row 6) (Fig. 2D) do not complement Fum1M strain sensitivity to HU. The expression levels of each of these constructs is similar relative to the loading control aconitase (Fig. 2E). The NES sequence once fused to FumA (FumA-NES) or FumB (FumB-NES), render these enzymes completely incapable of protecting Fum1M against DNA damage (SI Appendix, Fig. S2B, S2C). Figure S2D depicts expression of these modified proteins. Thus, localization of FumC to *the S. cerevisiae* nucleus is required for its DDR function.

### FumA and FumB enzymatic activity is required for their DNA damage response function

We have previously shown that the enzymatic activity of Class-II fumarases is required for their role in the DNA damage response [9, 12]. In order to ask whether the same is true regarding Class-I enzymes, FumA and FumB, we first needed to obtain enzymatically compromised mutants of these proteins. For this purpose, we designed substitution mutations, based on the predicted structure of *E. coli* Class-I fumarases, and according to the solved crystal structure of Class-I cytosolic FH of *Leishmania major* (5L2R, LmFH). We focused on the most conserved amino acids within a pocket/cavity that is suggested to be important for its catalytic activity [17]. Figure 3A depicts a dimeric homology model of the proposed structure for either FumA or FumB proteins, which share 90% amino acid identity and it is believed that they possess the same structure. The residues shown in spherical mode represent the dimeric enzyme interface that together form a cavity, that in LmFH forms the active site. The sticks represent conserved amino acids T236, I233 and Y481. Figure 3B shows alignment of the proposed *E. coli* Class-I fumarase structure (green) with that of *L. major* (blue, red, orange and yellow). We cloned PCR products harboring the point mutations into appropriate *E. coli* expression vectors. These plasmids were transformed into *E. coli* strains lacking each of the respective genes (Fig. 3C, 3D) and their expression was examined by Western blot (3E). Upon exposure to DNA damage (0.35% [v/v] MMS, 45 min), we observed an inability of the variant enzymes to protect against DNA damage in *E. coli* (Fig. 3C and 3D, compare rows 4 and 5 to row 3). The fumarase null mutant strain harboring these plasmids exhibits a highly significant reduction in enzymatic activity compared to wild-type FumA and FumB (Fig. 3F, 3G, compare bars 3 and 4 to bars 2). These results indicate that the enzymatic activity of FumA and FumB is crucial for their DDR related functions in *E. coli*.

### Alpha-Ketoglutarate complements DNA damage sensitivity of *E. coli fum* null mutants

In *S. cerevisiae* and human cells, the lack of extra mitochondrial fumarase and resulting sensitivity to DNA damage, can be complemented by externally added fumarate (fumaric acid, in the form of an ester, mono-ethyl fumarate, which is cleaved in the cells to form the free acid) [9, 10]. In *B. subtilis* the sensitivity of a fumarase null mutant to DNA damage, can be complemented by L-malate [12]. Both fumarate and L-malate are fumarase related enzymatic substrates/products. To examine if fumarase associated metabolites (substrates or products) complement the lack of fumarase in the DDR of *E. coli*, bacteria were grown in the presence of 25mM TCA cycle organic acids, added to the medium. An amazing result, as shown in Figure 4, is that *E. coli* strains deleted for the *fumA, fumB* and all three *fum* genes are protected from MMS induced DNA damage (0.35% [v/v] MMS, 45 min) <u>by α-KG</u> (bottom panel 2, compare rows 2, 3 and 6 to rows 1 and 4) and not by fumarate or L-malate (bottom panels 5 and 6 respectively). This complementation can be achieved by concentrations as low as 1mM α-KG added to the growth medium (SI Appendix, Fig. S3A). Growth of these strains on LB metabolite plates without MMS treatment, shows growth similar to that of the control (top panels). Even more surprising, is the fact that α-KG does not complement the DNA damage sensitivity of *fum* null mutants following irradiation (SI Appendix, Fig. S3B), demonstrating specificity to MMS treatment.

**Figure 4.**
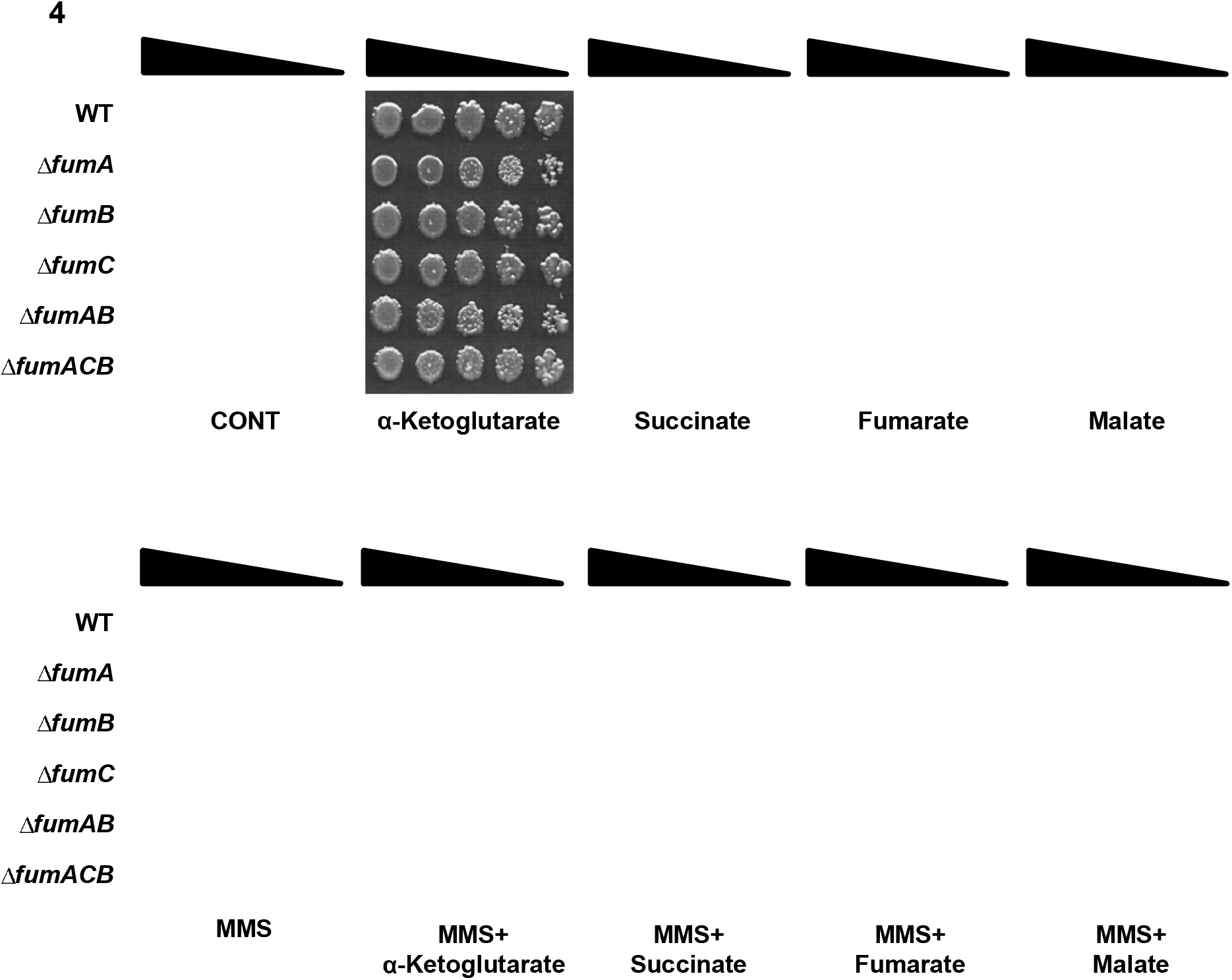
α-KG metabolically signal DNA repair in *fum* null mutant strains. *E. coli* wild type (WT), *ΔfumA, ΔfumB, ΔfumC, ΔfumAB* and *ΔfumACB* strains were grown to mid-exponential phase (OD600nm=0.3) and treated with MMS (0.35% MMS (v/v) for 45 min). The cells were serially diluted (spot test) onto LB plates containing or lacking the indicated organic acids. Each result is representative of three independent experiments.

This α-KG complementation of DNA damage sensitivity is exclusive to *fum* null mutants, as it does not protect *E. coli* strains harboring deletions in other DNA damage response genes or composite mutants of DDR genes and *fumA/fumB* (Fig S3C-E) from DNA damage induction, suggesting a direct relationship between fumarase and α-KG within the DNA damage response of *E. coli*.

To investigate this relationship, we decided to study the α-KG effect on two different levels; the first being to investigate a global impact of α-KG on transcription and induction of DNA repair pathways in *E. coli*. The second was to investigate a direct impact on the activity of DNA repair related enzymes.

### Absence of *E. coli* fumarases affects the transcript levels of many genes including Adaptive and SOS response genes <u>but does not involve α-KG signaling</u>

In *E. coli*, there are multiple DNA repair pathways, including; LexA-dependent SOS response triggering Homologous Recombination Repair (HRR), LexA-independent DNA damage responses and the Adaptive Response. MMS as well as UV-irradiation and other chemicals are capable of inducing the SOS response. The SOS response triggers HRR in *E. coli*, which proceeds via two genetic pathways, RecFOR and RecBCD. The RecBCD pathway is essential for the repair of DSBs and the RecFOR pathway for GAP repair [18]. The Adaptive Response involves the transcriptional induction of several genes in response to alkylation damage to the DNA caused by MMS treatment, among them *ada* (transcriptional regulator of the Adaptive response), *alkA, alkB*, and *aidB* [19, 20]. Given that the *fumA* and *fumB* single null and the *fumACB* triple null mutants are sensitive to DNA damage, compared to the wild type, and the apparent complementation of the DNA damage sensitivity phenotype by α-KG, we decided to study the transcriptional profiles of *fum* null mutants vs the WT strain via RNA-seq. The objective of the RNA-seq analysis was to find differentially expressed genes (DEGs, up or down regulated) in the WT strain in response to MMS treatment (0.35% [v/v],30 min), and more importantly in response to MMS+α-KG (0.35% [v/v], 30 min +25mM α-KG when indicated), compared to the mutant strains. This analysis was conducted in an effort to identify α-KG dependent enzymatic or regulatory components involved in the DNA damage sensitivity phenotype. Total RNA was extracted from three independent replicates, and RNA-seq analysis was performed (see materials and methods). To determine statistically significantly up or down regulated genes, a criterion of log ≥2.0-fold change or greater was used. The results are presented as Venn diagrams in figure 5A, 5B, S4A and S4B and Column diagram in Fig 5C. The corresponding raw data were deposited in NCBI GEO (accession: GSE161708). Figure 5A, 5B and S4A depicts changes in gene expression between the WT strain and the *fum* null mutant strains, following treatment with MMS or MMS + α-KG. Treatment with MMS coupled with α-KG does not significantly change the total number of genes with altered expression (S4B). All transcripts, shared and exclusive to each strain, were analyzed using the web Gene Ontology Resource tool [21, 22]. Of the upregulated genes, common to all tested strains (WT, *ΔfumA, ΔfumB* and *ΔfumACB*) we detect an upregulation of DNA repair and DNA recombination genes (SOS response genes, HRR genes, Adaptive response genes) and other stress response genes (oxidative stress response genes, various chaperones, and others). Of the downregulated genes, common to all tested strains (WT, *ΔfumA, ΔfumB* and *ΔfumACB*) we detect, downregulation of genes regulating cell growth, genes regulating cellular division and others. <u>Importantly, this is regardless of treatment with α-KG</u> in addition to MMS or not.

**Figure 5.**
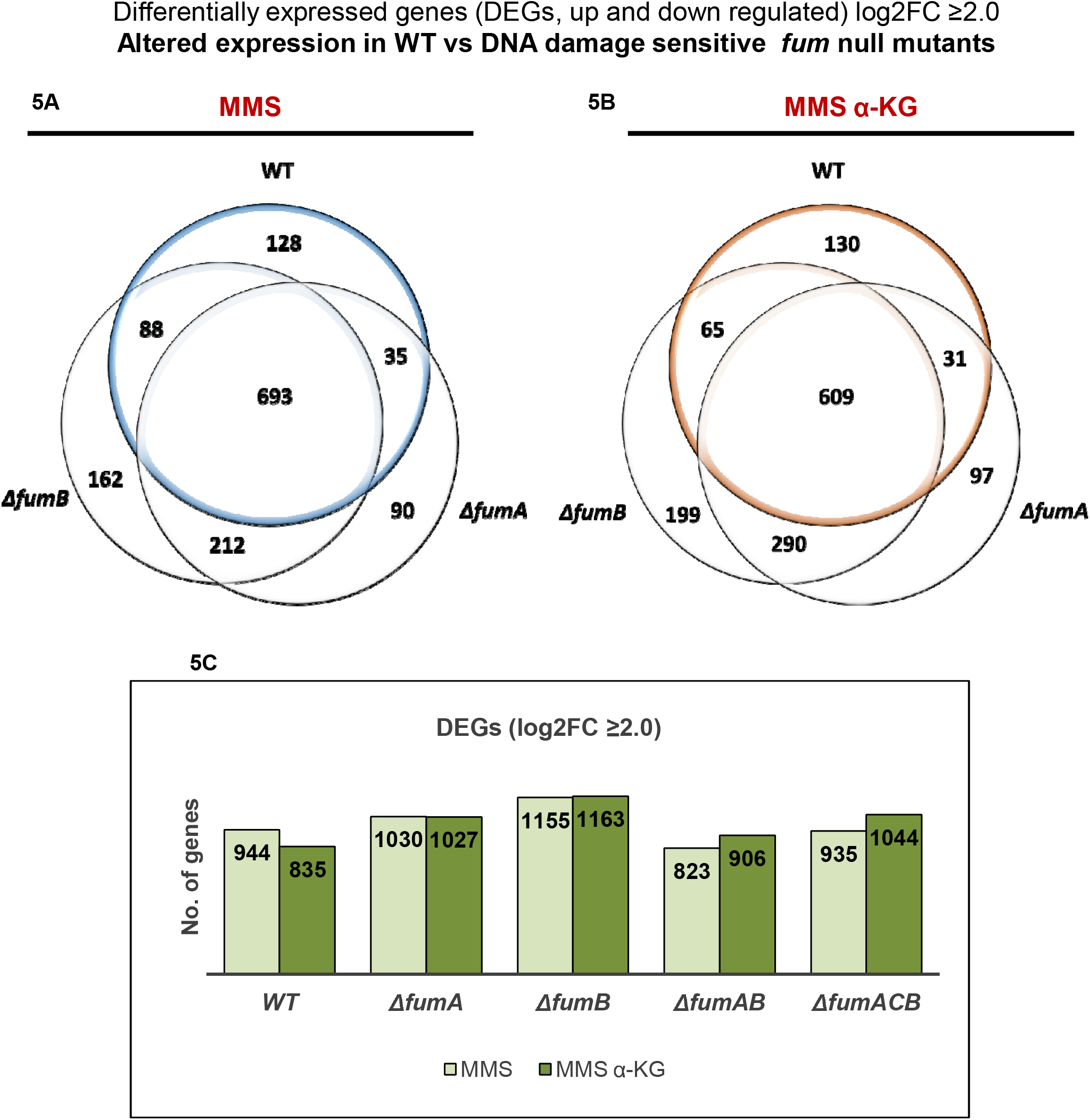
Absence of fumarase alters transcription in *E. coli*. RNA-seq analysis; *E. coli* wild type (WT), *ΔfumA, ΔfumB, ΔfumAB* and *ΔfumACB* strains were grown to early-exponential phase (OD600nm=0.3), and treated with MMS (0.35% MMS (v/v) for 30 min). The cells were collected and total RNA was extracted and subjected to RNA-seq analysis (see materials and methods). **(A, B)** Venn diagrams showing the common and differential genes (transcripts) between *E. coli* wild type (WT), *ΔfumA* and *ΔfumB* strains following treatment with MMS <u>or</u> MMS+25mM α-KG. **(C)** Column diagram (histogram) showing the number of differentially expressed genes in *E. coli* wild type (WT), *ΔfumA, ΔfumB, ΔfumAB* and *ΔfumACB* strains following treatment with MMS <u>or</u> MMS+25mM α-KG.

Most importantly, analyzing DEGs altered exclusively in the WT strain or exclusively in either *fum* mutant strain (supplementary table 1), yielded no candidate genes that may explain the DNA damage sensitivity of *ΔfumA, ΔfumB* and *ΔfumACB* strains, as all DNA repair systems seem to be induced in all strains and conditions. For this reason, we decided to look specifically at the induction levels (fold change in expression) of DNA repair genes and see if there is any direct effect resulting from the absence of fumarase. Examples of SOS response genes (*recA, umuD, dinI* and *ssb*), Adaptive response genes (*aidB, alkA* and *alkB*) and genes controlling cell division (*sulA, ftsZ, minC* and *minD*) and cellular chaperone (*dnaK*) are presented in Figure S4C-N. As mentioned above, we find induction of the SOS and the Adaptive response genes in the presence of MMS or MMS+ α-KG in all strains, coupled with the decrease in expression of cell division genes, demonstrating a need for inhibition of the cell cycle for the repair of the damaged DNA. For *recA, umuD, aidB, alkA, alkB, ssb* there is a significantly higher expression level in the WT strain compared to the *fum* null mutant strains following MMS treatment. It is important to emphasize again that <u>α-KG has no additive effect to treatment with MMS alone</u> on the expression levels of these genes. In this regard, we find insignificant differences in the levels of α-KG in the extracts of wild type (WT), *ΔfumA, ΔfumB* and *ΔfumC* cells, in the presence or absence of MMS (by LC-MS). Thus, the cellular levels of α-KG cannot explain this phenomenon.

In order to validate the reproducibility and accuracy of the RNA-seq analysis results, we determined the mRNA levels of four DDR and stress response genes by RT-qPCR; *recA, dinI, alkA* and *alkB*. All strains were grown as per the RNA-seq experiment in three independent replicates (0.35% [v/v] MMS, 30 min +25 mM α-KG when indicated). The results are consistent with those of the RNA-seq data (SI Appendix, Fig. S5A-D). Primers used for this assay are presented in the material and methods section.

Why do *E. coli fum* null mutants exhibit a lower expression of these various DDR genes, or a higher expression level of others g? Whether this effect is mediated by fumarase metabolites, fumarate or malate, as seen in other systems remains to be studied [23–25].

### Alpha-ketoglutarate enhances, whereas succinate and fumarate inhibit AlkB DNA damage repair activity

Our most important conclusion from the previous section is that the candidate DNA damage signaling metabolite, α-KG, does not function at the level of transcription. Given these results, we chose to focus on DNA repair enzymes whose activity may be regulated by TCA cycle organic acids. We decided to focus on AlkB due to its known function in DNA damage repair and its dependency on α-KG. AlkB is a Fe-(II)/α-KG-dependent dioxygenase in *E. coli*, which catalyzes the direct reversal of alkylation damage to DNA, primarily 1-methyladenine (1meA) and 3-methylcytosine (3meC) lesions. A hallmark of AlkB-mediated oxidative demethylation is that the oxidized methyl group is removed as formaldehyde [26, 27]. Previously published research in the bacterium *B. subtilis* showed that in fumarase null mutants, there is an accumulation of succinate and fumarate [12]. In eukaryotes, fumarate and succinate have been shown to inhibit human α-KG-dependent histone and DNA demethylases and in yeast, fumarate inhibits the sensitivity of components of the homologous recombination repair system to DNA damage induction [10, 11, 28]. The inhibition of histone and DNA α-KG-dependent demethylases by fumarate and succinate was established as competitive, based on their structural similarity to α-KG [28].

To test whether fumarate and succinate inhibit AlkB demethylase activity in *E. coli*, we employed two previously published protocols [29, 30] for in vitro measurement of DNA damage repair by AlkB in the presence and absence of succinate, fumarate and malate. Such an effect, if found could explain the increase in sensitivity of *fum* null mutants to MMS generated DNA damage, as the absence of fumarase leads to accumulation of fumarate and succinate.

The first approach was direct fluorescence-based formaldehyde detection using acetoacetanilide and ammonium acetate, which together form an enamine-type intermediate. This intermediate undergoes cyclodehydration to generate highly fluorescent dihydropyridine derivative, having maximum excitation at 365 nm and maximum emission at 465 nm [30]. A 70bp oligonucleotide (see materials and methods) was treated with MMS to allow methylation/alkylation, and was directly incubated with purified AlkB (2.5μM) to facilitate DNA repair. DNA repair was measured by release of formaldehyde (RFU [HCHO concentration]). While α-KG is required for the full activity of AlkB in vitro (Figure 6A, bar 1, 200μM of α-KG), succinate and fumarate competitively inhibit *E. coli* AlkB demethylase activity in a dose dependent manner (compare bars 2-4, 5-7 to bar 1) (200μM of succinate and fumarate with increasing amounts of α-KG). Malate appears to have a small insignificant, non-dose dependent, effect on AlkB demethylase activity (bars 8-10) (200μM of malate with increasing amounts of α-KG). The second approach, employs a restriction enzyme-based demethylation assay [29]. The same substrate (70bp oligonucleotide substrate that contains the dam sensitive, MboI restriction recognition site in the middle of the sequence) was treated with MMS to allow methylation, and was directly incubated with purified AlkB (Fig. 6B), or *E. coli* WT and *fum* null mutant strains whole cellular lysate (treated with MMS to induce AlkB expression) (SI Appendix, Fig. S6A,B) to facilitate DNA repair. By digestion with the dam methylation sensitive enzyme MboI, successful repair of methylated oligonucleotide (complete demethylation by AlkB) yields a 35bp product (6B, compare lane 4 to lane 3 [methylated dsDNA [me-dsDNA] undigested by MboI] and lane 2 [control un-methylated dsDNA digested by MboI]). Shown in Figure 6B, is the inhibition of AlkB demethylase activity which can be detected by the non-apparent 35bp fragment. Succinate and fumarate inhibit AlkB in a dose dependent manner (compare lanes 5-8 [containing 200 μM succinic acid and increasing concentrations of α-KG] and lanes 9-12 [200 μM fumaric acid and increasing concentrations of α-KG] to lane 4 [containing 200 μM α-KG]). Malate appears to have an insignificant effect on AlkB demethylase activity (lanes 13-16). Together, these results demonstrate, for the first time, *E. coli* AlkB inhibition by fumarate and succinate. Figure S6 depicts an in vitro AlkB activity assay followed by MboI digestion, using whole cellular lysate from the WT and *fum* null mutant strains, in which all strains exhibit repair of methylated ssDNA.

**Figure 6.**
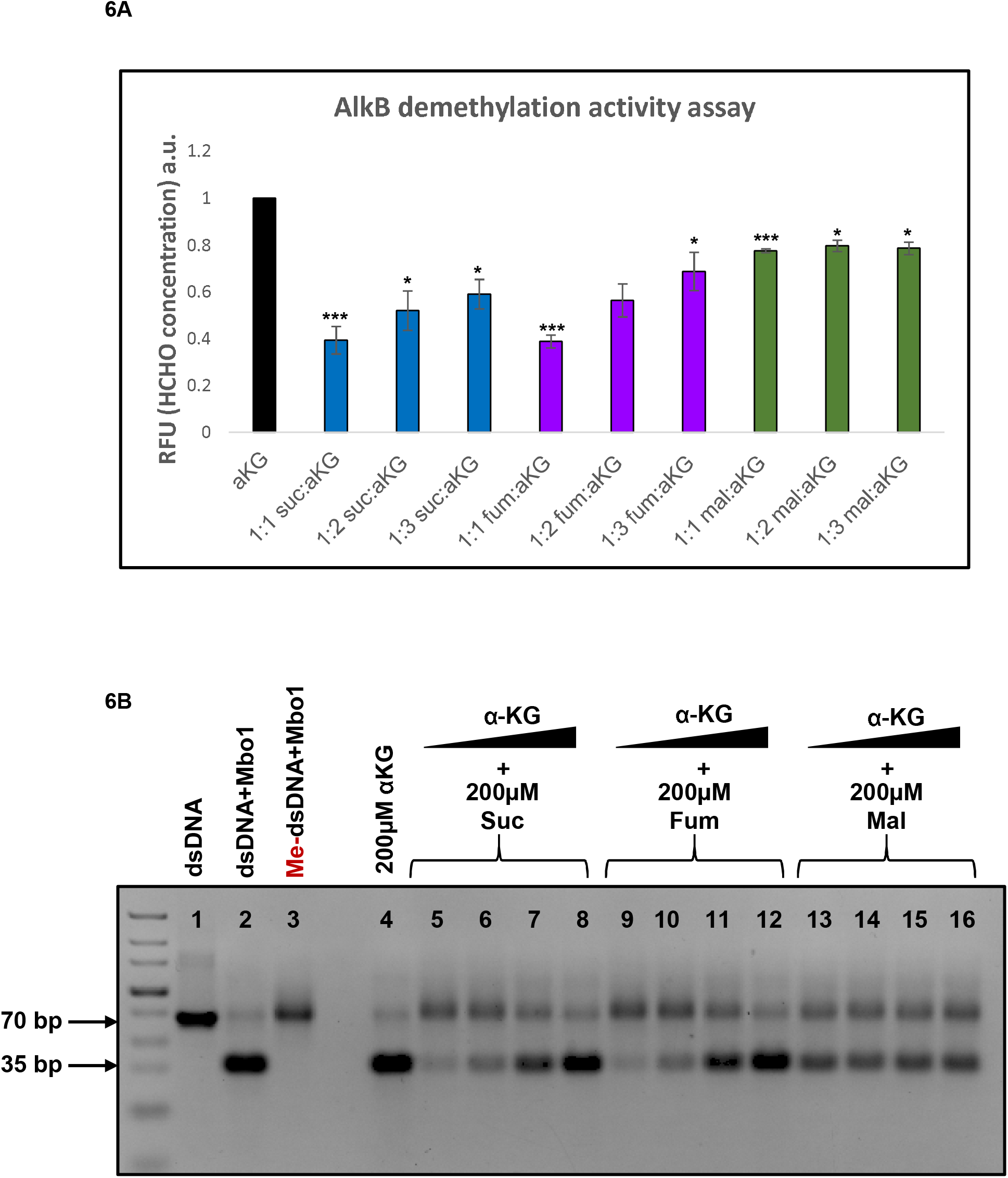
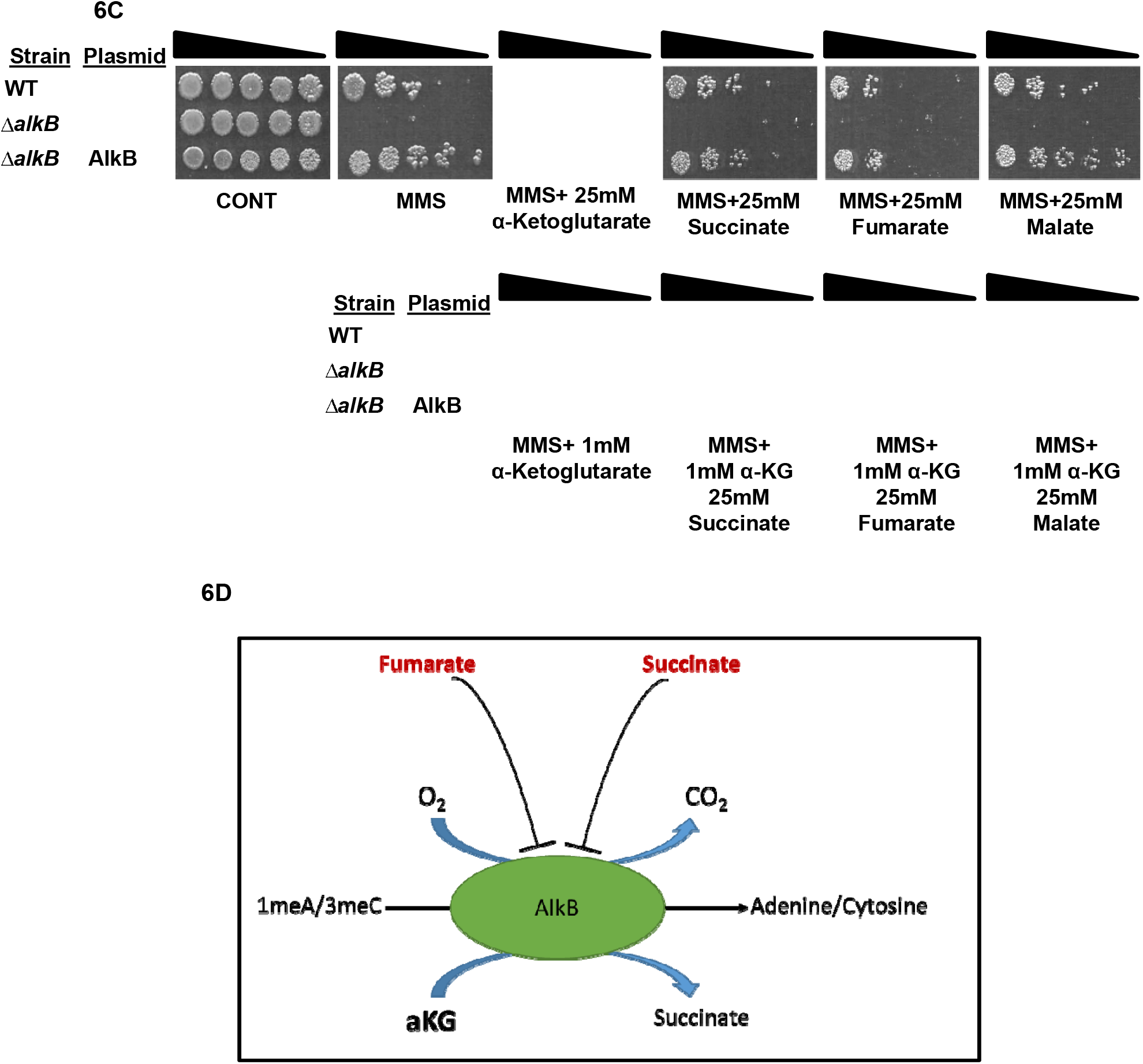
Fumarate and succinate inhibit AlkB demethylase activity. (**A**) Repair of methylated oligonucleotides for 1 hour at 37□C by 2.5μM AlkB. DNA repair was quantified by measuring formaldehyde release. The repair reaction was mixed following in vitro repair, with ammonia and acetoacetanilide, incubated at room temp for 15 minutes, forming a highly fluorescent dihydropyridine derivative. The reaction was analyzed using a multimode reader setting the excitation wavelength at 365 nm and emission wavelength at 465 nm. The graph is represented as mean ± standard error (*n=3*, two-tailed *t*-test *p < 0.05; **p < 0.01; ***p < 0.005). (**B**) Repair of methylated oligonucleotides for 4 hours at 37□C by 2.5μM AlkB. Following in vitro repair, the reaction was mixed with equi-molar concentration of complementary ssDNA, the resultant dsDNA was digested with MboI and subjected to agarose electrophoresis and staining with safeU. Following successful demethylation, digestion with MboI produces a 35bp product. (**C***) E. coli* wild type (WT), *ΔalkB* and *ΔalkB* expressing AlkB were grown to mid-exponential phase (OD600nm=0.3) and treated with MMS (0.35% MMS (v/v) for 45 min). The cells were then washed and serially diluted (spot test) onto LB plates and LB plates containing the indicated organic acids. **(D)** Proposed model for the participation of α-KG, fumarate and succinate in the regulation of AlkB activity, in DNA damage sensitive *fum* null mutants in response to MMS treatment.

In order to validate these results in vivo, an AlkB inhibition experiment with high molar concentrations (25mM) of fumarate and succinate was carried out. For this purpose, we induced DNA damage (using 0.35% [v/v] MMS, 45 min) in an *E. coli* strain lacking *alkB* (*ΔalkB*), and in a *ΔalkB* strain, overexpressing AlkB from a plasmid. Figure 6C shows that *ΔalkB* is extremely sensitive to MMS generated DNA damage, and overexpression of AlkB completely compensates for this DNA damage sensitivity phenotype (panel 2, compare row 3 to row 2). Fumarate and succinate exhibit a strong inhibition of AlkB ability to repair damaged DNA (compare top row, panels 4 and 5 to panels 2 and 3). Addition of as low as 1 mM of α-KG, in addition to succinate or fumarate complements AlkB activity in vivo, and alleviates the fumarate and succinate growth inhibition (compare bottom row, panels 2 and 3 to top row panels 4 and 5). These results are consistent with the previously shown in vitro experiments above, in which fumarate and succinate strongly inhibit AlkB DNA repair activity with malate having no effect on the AlkB DNA repair function. These results are summarized in a model in which accumulation of fumarate and succinate competitively inhibit the α-KG-dependent-AlkB activity, leading to failure of DNA repair and subsequently cellular death, explaining the sensitivity of *ΔfumA, ΔfumB* and *ΔfumACB* to MMS induced DNA damage (Fig. 6D).

## Discussion

Recent discoveries identify the recruitment of metabolic intermediates and their associated pathways in the signaling of crucial cellular functions. The best example is the enzyme fumarase (fumarate hydratase) which belongs to Class-II (FumC-like fumarases). This TCA cycle enzyme has been shown to be involved in the DNA damage response in yeast and human, and more recently in the Gram-positive bacterium *B. subtilis* [10–12, 31]. In human and yeast, fumarate (fumaric acid) is the signaling molecule targeting histone demethylases and a HRR resection enzyme respectively [10, 11]. In *B. subtilis* L-malate is the signaling molecule, modulating RecN (the first enzyme recruited to DNA damage sites) expression levels and cellular localization [12].

In this study we show an amazing variation on the above themes by studying *E. coli*, which harbors the two Classes of fumarases, Class-I *fumA* and *fumB* and Class-II *fumC*. At first glance, contrary to previous work on Class-II fumarases, *E. coli* FumC does not have a natural role in the DDR, while FumA and FumB, harboring no structural or sequence similarity to the Class-II fumarases, are participants in the DDR in this Gramnegative bacterium. In this regard, not only do FumA and FumB participate in the DDR in *E. coli*, they can substitute for yeast fumarase in a Fum1M DNA damage sensitive model strain [9, 12]. This analysis shows, that Class-I fumarases, FumA/FumB, that are non-homologous to the yeast Class-II fumarase, can function in the DDR, indicating that not the enzyme is needed for this DDR related function, but rather its enzymatic activity and related metabolites. As we have demonstrated for Class-II FH in human cell lines, yeast and *B. subtilis*, Class-I FumA and FumB enzymatic activity is crucial for their DNA damage protective function. This is supported in this study by the fact that enzymatically compromised variants of the Class-I fumarase do not complement DNA damage sensitivity. Our results indicate that it is only the level of fumarase activity that is important, since over-expression of all *E. coli fum* genes can function in the DDR.

Most interesting is, that neither fumarate nor malate added to the growth medium are capable of complementing the MMS induced DNA damage sensitive phenotype of *fum* null mutants as demonstrated for the previously studied organisms. Rather, α-KG, a TCA cycle intermediate with no direct interaction with fumarase appears to be the signaling metabolite for this adaptive response related damage. Moreover, this α-KG mediated protective effect seems to be exclusive to *fum* null mutants, as the sensitivity phenotype of other *E. coli* null mutants in DNA damage repair genes, is not alleviated by the addition of α-KG to the growth medium. Fumarase absence, and perhaps alteration of local concentration of TCA cycle metabolites, specifically affects the activity of the DNA repair enzyme AlkB, on the one hand, and leads to a reduced transcriptional induction of DNA damage repair components, on the other.

The absence of fumarase is shown here to affect AlkB, a Fe-(II)/α-KG-dependent dioxygenase via its precursor metabolites succinate and fumarate. Fe-(II)/α-KG-dependent dioxygenases are present in all living organisms and catalyze hydroxylation reactions on a diverse set of substrates, including proteins, alkylated DNA/RNA, lipids and others [32, 33]. Fumarate and its precursor succinate have been shown to inhibit a variety of Fe-(II)/α-KG-dependent dioxygenases, and this inhibition was shown to antagonize α-KG-dependent processes and negatively regulate Fe-(II)/α-KG-dependent dioxygenases such as prolyl hydroxylases (PHDs), histone lysine demethylases (KDMs), collagen prolyl-4-hydroxylases, and the TET (ten-eleven translocation) family of 5-methlycytosine (5mC) hydroxylases [10, 28, 31, 34–37]. We find, in agreement with the above observations, that AlkB enzymatic activity is inhibited by the presence of high concentrations of fumarate and succinate both in vitro and in vivo, indicating that AlkB inhibition has a specific consequence in vivo, leading to the sensitivity of *ΔfumA, ΔfumB* and *ΔfumACB* to MMS induced DNA damage.

The above findings suggest that the recruitment of metabolic intermediates and their associated pathways in signaling during evolution, was adaptable. In an ongoing study we are examining the distribution of Class-I and Class-II fumarases through the different domains of life. Class-I and Class-II can be found in prokaryotes and eukaryotes as sole fumarase representatives; e.g., mammals harbor only Class-II while many protozoa contain only Class-I fumarases. Some organisms, such as *E. coli*, harbor both Classes and, in fact, very few organisms can be found that lack both Classes of fumarases. At this stage it is difficult for us to hypothesize why a certain metabolite, Class of enzyme, or pathway was chosen to signal the DDR, but this study gives us a new perception on how metabolites, metabolic enzymes and pathways could have been recruited during evolution of metabolite signaling.

There are many unanswered questions regarding the fumarases in *E. coli*, how fumarase and fumarate regulate different types of DNA damage repair, and in particular, why *ΔfumAB* behaves differently than *ΔfumA* and *ΔfumB* strains? The increase in FumC and its participation in the DDR is one possible mechanism and a second may be the localization of the FumC to different DNA associated proteins or subcellular locations. In this regard, using specific Class-I and Class-II antibodies we detect that the enzymes do not co-localize with the nucleoid but rather localize to the cell poles regardless of treatment with MMS, as seen for other DNA damage repair enzymes [38]. Another explanation is that FumA and FumB are able to form a heterodimer for which we have very preliminary data (IP, Co-IP and Mass Spec analysis). What governs the levels of α-KG, fumarate and other metabolites at sites of DNA damage? And how exactly does fumarate exert its effect on transcription? We have only examined the facultative anaerobe *E. coli* under aerobic conditions, and the question is whether the metabolic regulation of the DDR that we have uncovered, changes under anaerobic growth conditions. Further studies are needed to address these complex questions and systems.

## Materials and Methods

### 1. Bacterial Strains and Growth Conditions

Bacterial strains were provided by the Coli Genetic Stock Center and are listed in supplementary table 2. Additional strains were constructed using the lambda red system [39], plasmids were constructed using standard methods, plasmids and primers are listed in supplementary table 3. Bacteria were grown in Luria-Bertani (LB) broth to early-mid-exponential phase (OD_600nm_=0.30), supplemented, when needed, with ampicillin (Amp, 100 g/ml) or 25 mM metabolites (α-KG, succinate, fumarate, malate [Sigma Aldrich]). MMS was added to a final concentration of 0.35% [v/v]. When indicated 0.5 mM of IPTG (isopropyl-D-thiogalactopyranoside, Sigma) was added for expression of genes from the Puc18 vector under the Lac promoter. For agar plates, 2% agar was added.

### 2. Yeast Strains and Growth Conditions

*S. cerevisiae* strains used in this research are listed in supplementary table 4. *S. cerevisiae* strains used were grown on synthetic complete (SC) medium containing 0.67% (wt/vol) yeast nitrogen base +2% glucose (SC-Dex) or 2% galactose (SC-Gal) (wt/vol), supplemented with appropriate amino acids. For agar plates, 2% agar was added. HU (hydroxyurea) was added to SC medium (supplemented with glucose) to a final concentration of 200mM. The cells were grown overnight at 30°C in SC-Dex, washed in sterile double distilled water, and transferred to SC-Gal for induction of genes under GAL promoter.

## 3. Construction of mutants- Gene knockout using λ red system (pKD46)

### 3.1. Gene Disruption

Δ*fumAB*, Δ*fumACB* strains (and other composite mutants in this study) were constructed by the following: Gene disruption as previously described [39]. Essentially, Δ*fumB* and BW25113 Transformants carrying λ red plasmid were grown in 5-ml LB cultures with ampicillin and L-arabinose at 30°C to an OD_600_ of 0.6. The cells were made electrocompetent by concentrating 100-fold and washing three times with ice-cold sterile DDW. Electroporation was done by using a Cell-Porator with a voltage booster and 0.2-cm chambers according to the manufacturer’s instructions (BIO-RAD). By using 100 μl of suspended cells and 100 ng of PCR product (for Δ*fumAB* strain cat [Chloramphenicol, Cm] cassette with flanking upstream and downstream sequences of *fumA*, and kan cassette [Kanamycin] with flanking upstream and downstream sequences of *fumB*, and for Δ*fumAC* strain, cat cassette with flanking upstream of *fumA* and downstream of *fumC*), the cells were added to 1ml LB, incubated 3 hours at 37°C, and then spread onto agar supplemented with Cm/Kan to select CmR/KanR transformants. After primary selection, mutants were colony purified twice selectively at 37°C and then tested for loss of the parental gene by PCR.

Following verification of *fumAC* knock out and cassette insertion, a second Knock out was carried out with Kan (Kanamycin) cassette with flanking upstream and downstream of *fumB* [two step Gene knockout]) resulting in the Δ*fumACB* strain.

### 3.2. PCR verification of gene disruption

Two PCR reactions were used to show that the mutants lost the respective *fum* genes. A freshly isolated colony was used in separate PCRs following a 5-min pre-incubation ‘‘hot start,” at 95°C. One reaction was done by using nearby locus-specific primers (150bp upstream of *fumA* or *fumB*) with the respective common test primer (c1 and k2) to verify gain of the new mutant specific fragment (antibiotic resistance), and a second reaction was carried out with the flanking locus-specific primers to verify simultaneous loss of the parental (non-mutant) fragment. Control colonies were always tested side-by-side.

## 4. Prediction of FumA and FumB structure

Prediction of FumA and FumB were as previously described [40]. Essentially, Homology models of the structure of *E. coli* FumA and FumB (based on the solved structures of 5L2R of *L. major*, [17]) were generated using I-TASSER [41]. The models with the highest probability score among the provided models were selected for further investigation. Using PeptiMap [42] for the identification of peptide binding sites and CONSURF [43] for conservation analysis of FumA and FumB amino acid sequence. Mutations were designed based on conserved amino acids within the peptide binding pockets detected. Figures of structures were generated using the PyMOL software (Schrödinger Inc).

### 4.1. Mutagenesis of suspected catalytic site

Three proposed amino acids suspected to be vital for catalytic activity of FumA and FumB; Isoleucine 233, Threonine 236 and Tyrosine 481 were mutagenized by site directed mutagenesis using a KAPA HiFi HotStart PCR kit (KAPABIOSYSTEMS). Exchanging Tyrosine, at 481 with Aspartic acid (Y481D), we chose to combine I233 and T236 generating a double point mutation, exchanging Isoleucine at 233 for lysine, and exchanging Threonine at 236 with Isoleucine (I233K T236I). All mutated genes were cloned into Puc18 vector and transformed into bacterial strains Δ*fumA*, Δ*fumB* and Δ*fumACB*.

## 5. FumA/FumB activity assay

FumA and FumB activity were assayed as previously described [13]. Essentially, fumarase enzymatic activity was measured by monitoring malate depletion at 250 nm spectrophotometrically in quartz cuvettes (Fumarase activity at 250 nm with L-malic acid as the substrate). Assays were performed in assay buffer (100 mM Hepes buffer pH 8 with 5% glycerol and 0.5 mM DTT).

## 6. RNA-seq analysis

### 6.1. Samples preparation, RNA extraction and library preparation

Bacterial pellet samples (untreated and treated- 0.35% [v/v] MMS, 30 min with and without α-KG) were submitted to Axil Scientific NGS, SG. Total RNA extraction was performed using the total RNA purification kit from Norgen Biotek Corp. The quality and quantity of RNA were measured using Nanodrop and Tapestation 4200. Library was prepped using Ribo-zero bacteria and mRNA library prep kit according to the manufacturer’s instructions.

### 6.2. Sequencing and analysis

Next, the libraries were sequenced on NextSeq 500 machine. Raw sequencing data were received in FASTQ formatted file and are deposited in NCBI GEO (accession number GSE161708). Quality control steps were performed using fastp v0.19.4 [44] to access bases quality of each sequences and also to filter low quality bases (Phred 33 quality < Q20) and adapter sequences. By using BWA v7.6 [45] tool, quality filtered sequences were then mapped to *Escherichia coli* BW251153 (WT) reference genome (accession number: CP009273) obtained from GenBank.

The RNA-seq generated 6.6-104.9 million raw reads with a remaining 5.8-92. 5 million clean reads that were subsequently mapped to *E. coli* genome. 5.8-91.3 million were mapped with a mapped fraction of 94.4-100% (Supplementary table 5). Resulting alignment files of each samples were used in downstream analysis of differential expression. The mRNA expression was then quantified using StringTie [46] tool. Three expression values were computed which are reads count, FPKM (fragments per kilobase million) and TPM (transcripts per kilobase million). Differential expression (DE) analysis was performed using DESeq2 tool to compare expression levels of all mRNA under different experimental conditions and the expression fold change between each comparison group was calculated and log-transformed. mRNA with expression fold change ≥2.0 and p-adjusted <0.01 were selected as statistically significant differentially expressed.

## 7. RT-qPCR analysis

Bacteria were grown and treated with MMS or MMS+ α-KG as in RNA-seq experiment. Total RNA was isolated using a zymo Quick-RNA Miniprep Kit (R1055). Samples of 1μg total RNA were reverse transcribed to cDNA using iScript^TM^ Reverse Transcription supermix for RT-qPCR (Bio-Rad 1708841) according to manufacturer’s instructions. cDNA samples were PCR amplified using iTaq^™^ Universal SYBR Green Supermix (Bio-Rad 1725121). Real-time quantitative PCR was carried out on the ABI 7500 Fast Real-Time PCR System (Applied Biosystems).

The following primers were used for cDNA amplification:

*alkA* 5’-GCCAGACTACGGGCATAATAAC-3’ (forward) and 5’-GGTCGTGGATGTTGGGATTT-3’ (reverse)

*alkB* 5’-CCAGCCAGATGCTTGTCTTATC-3’ (forward) and 5’-GCAGATCCGGTTCGTCTTTATC-3’ (reverse)

*recA* 5’-GGCTGAATTCCAGATCCTCTAC-3’ (forward) and 5’-CTTTACCCTGACCGATCTTCTC-3’ (reverse)

*dinI* 5’-TCGCGCCAATAACCGATAAA-3’ (forward) and 5’-GAACTTTCCCGCCGTATTCA-3’ (reverse)

*gyrA* 5’-TCAGCGGAGAACAGCATTAC-3’ (forward) and 5’-CCGGTAAAGTGGCGATCAA-3’ (reverse)

## 8. AlkB in vitro demethylation assay

AlkB demethylation activity was measured by two previously described methods [29, 30]. For both methods 40 μg of chemically synthesized oligonucleotides (IDT) that contains the restriction site for the methylation sensitive restriction enzyme MboI (underlined) was used 5’-GGATGCCTTC GACACCTAGC TTTGTTAGGT CTG<u>GATC</u>CTC GAAATACAAA GATTGTACTG AGAGTGCACC-3’). The oligonucleotides were treated with MMS buffer (5% (v/v) MMS, 50% EtoH in a final volume of 500 μl in presence of 200 mM K_2_HPO_4_ at room temperature for 16 hours), dialyzed in TE buffer (10 mM Tris pH 8.0, 1 mM EDTA pH 8.0 [Sigma Aldrich E5134-50G]) using Spectra/Por dialysis membrane (MWCO: 3500) and precipitated with 2 volumes of ice cold 100% EtoH and 0.3M sodium acetate pH 5.5, washed twice with 70% EtoH and dissolved in DNase/RNase free water.

### 8.1 Immunoprecipitation of FLAG tagged AlkB-

AlkB was expressed in *E. coli* BL21 and Immunoprecipitated using FLAGIPT1 kit (sigma Aldrich) according to manufacturer’s instructions, and Flag-AlkB proteins were eluted using a Flag peptide.

### 8.2 Direct fluorescence-based formaldehyde detection

This assay was performed as previously described [30]. Repair reactions (50 μl) were carried out at 37□C for 1 hour in the presence of 2.5 μM AlkB, 0.5 μg (1 μM) methylated oligonucleotide and reaction buffer (20 mM Tris-HCl pH 8.0, 200 μM indicated metabolite [αKG acid, succinic acid, fumaric acid and malic acid], 2 mM L-Ascorbate and 20 μM Fe(NH_4_)_2_(SO_4_)_2_). Formaldehyde release was detected by mixing demethylation repair reaction product with 40 μl of 5 M ammonium acetate and 10 μl of 0.5 M acetoacetanilide to make the final volume 100 μl. The fluorescent compound was allowed to develop at room temperature for 15 min and then entire reaction mixture was transferred to 96-well microplate and analysed using a multimode reader setting the excitation wavelength at 365 nm and emission wavelength at 465 nm. Formaldehyde standard curve was prepared by selecting a range of pure formaldehyde concentrations from 2 to 20 μM.

### 8.3 Restriction based demethylation assay

This assay was performed as previously described [29]. Repair reactions (50 μl) were carried out at 37□C for 4 hours in the presence of 2.5 μM AlkB, 0.5 μg (1 μM) methylated oligonucleotide and reaction buffer (20 mM Tris-HCl pH 8.0, 200 μM indicated metabolite, 2 mM L-Ascorbate and 20 μM Fe(NH4)2(SO4)2). Repair reaction was annealed to equi-molar complimentary ssDNA to generate methylated dsDNA in annealing buffer (50 mM Hepes pH 8 and 10 mM EDTA pH 8) at 37°C for 60 min. repaired dsDNA was digested by MboI restriction enzyme (2 hours at 37°C followed by heat inactivation). Digestion products were dissolved on 3% agarose containing SafeU (SYBR^™^ Safe DNA Gel Stain, Thermo Fisher Scientific, #S33102) with 10 mM sodium borate as electrophoresis buffer at 300 V for 20 min, and visualized using the Gel Documentation System (Bio Rad).

## 9. Western blot analysis

*E. coli* cells were harvested in lysis buffer containing: 50 mM Tris pH 8, 10% glycerol, 0.1% triton, 100 mM PMSF (Sigma Aldrich 10837091001), 0.1 mg/ml Lysozyme (Sigma Aldrich 10837059001). Protein concentrations were determined using the Bradford method [47]. Loading amount on gels was normalized using protein concentration and verified by recording the gels using Ponceau S staining. Samples were separated on 10% SDS-PAGE gels, transferred onto PVDF membranes (Millipore). The following primary antibodies were used: polyclonal anti FumA, polyclonal anti FumC and for identification of Class-I FH polyclonal anti FumB that recognizes both FumA and FumB was used (prepared in our lab).

*S. Cerevisiae* cells were harvested in lysis buffer containing: 10 mM Tris pH 8, 1 mM EDTA and 100 mM PMSF. Protein concentrations were determined using the Bradford method [48]. Samples were separated on 10% SDS-PAGE gels, transferred onto PVDF membranes (Millipore). The following primary antibodies were used: monoclonal anti FLAG (Sigma- F3165) and polyclonal anti Aconitase (prepared in our lab). All blots were incubated with the appropriate IgG-HRP-conjugated secondary antibody. Protein bands were visualized and developed using ECL immunoblotting detection system (ImageQuant LAS 4000 mini GE Healthcare) and Gel Documentation System (Bio Rad).

## 10. Statistical analysis

Statistical analysis was conducted with the two-tailed unpaired Student’s t-test. All data represent the mean ± standard error of three independent experiments.

## Author Contributions

Yardena Silas, Conceptualization, Resources, Validation, Investigation, Visualization, Methodology, Writing—original draft, Writing—review and editing; Esti Singer, Validation, Investigation, Visualization, Methodology, Writing—review and editing; Norbert Lehming Methodology, review and editing; Ophry Pines, Conceptualization, Resources, Supervision, Funding acquisition, Validation, Writing—original draft, Project administration, Writing—review and editing.

## Supplemental information

**Figure S1.**
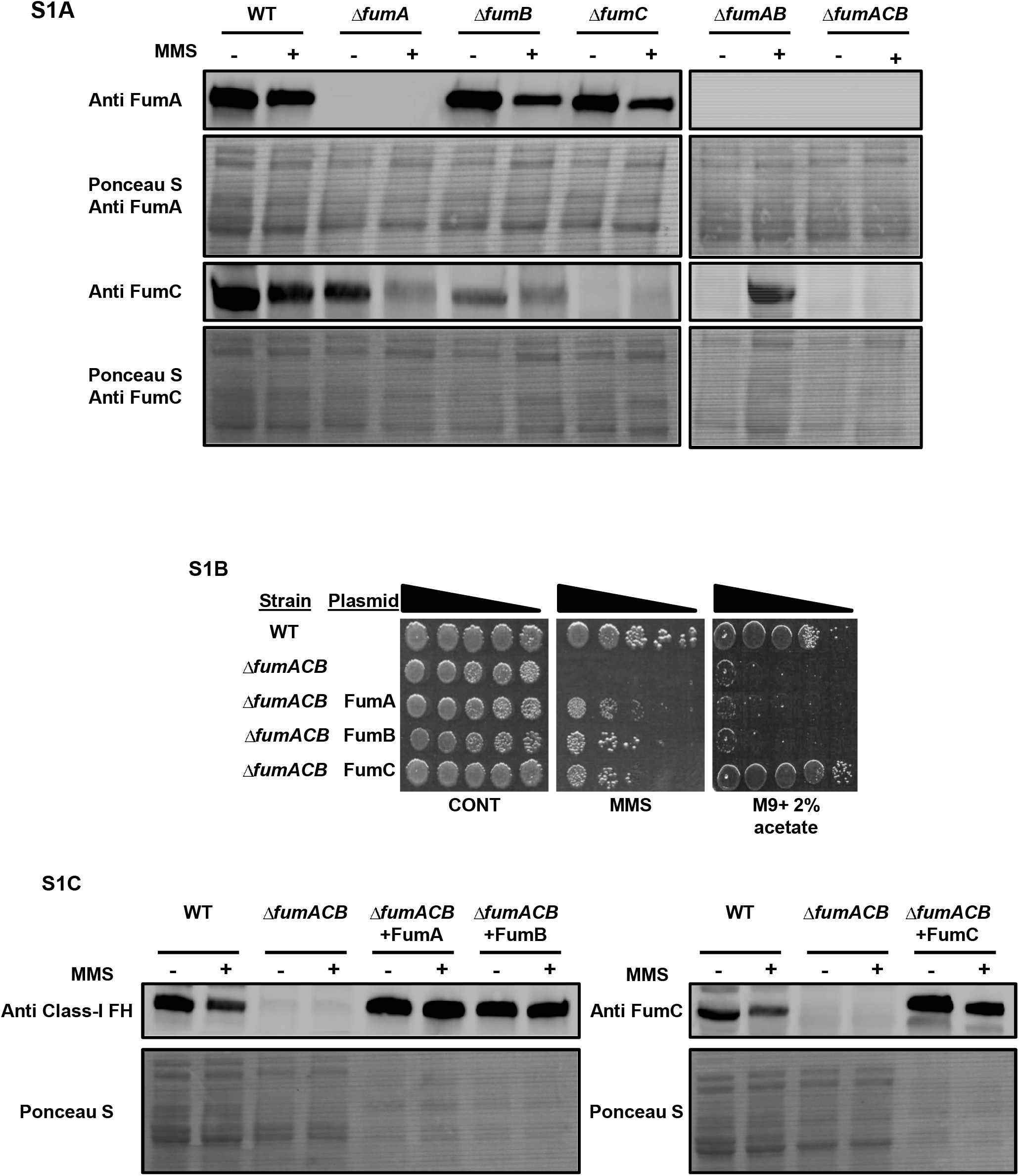
In the absence of FumA and FumB, FumC participates in the DDR in *E. coli*. (**A**) *E. coli* wild type (WT), *ΔfumA, ΔfumB, ΔfumC, ΔfumAB, ΔfumACB* strains were grown to mid-exponential phase and treated with MMS (0.35% (v/v) for 45 minutes at 37□C). Total cell extracts were prepared from the indicated samples and subjected to Western blotting, using the indicated antibodies followed by Ponceau S staining. Each result is representative of three independent experiments. (**B**) *E. coli* wild type (WT), *ΔfumACB* and *ΔfumACB* harboring plasmids encoding the indicated *E. coli fum* genes grown to early-exponential phase (OD600nm=0.3), and treated with MMS (0.35% [v/v] for 45 minutes at 37□C). The cells were then washed and serially diluted (spot test) onto LB plates. (**C**) *E. coli* wild type (WT), *ΔfumACB* and *ΔfumACB* harboring plasmids encoding the indicated *E. coli fum* genes were grown as in S1A. Total cell extracts were prepared from the indicated samples and subjected to Western blotting, using the indicated antibodies followed by Ponceau S staining. Each result is representative of three independent experiments.

**Figure S2.**
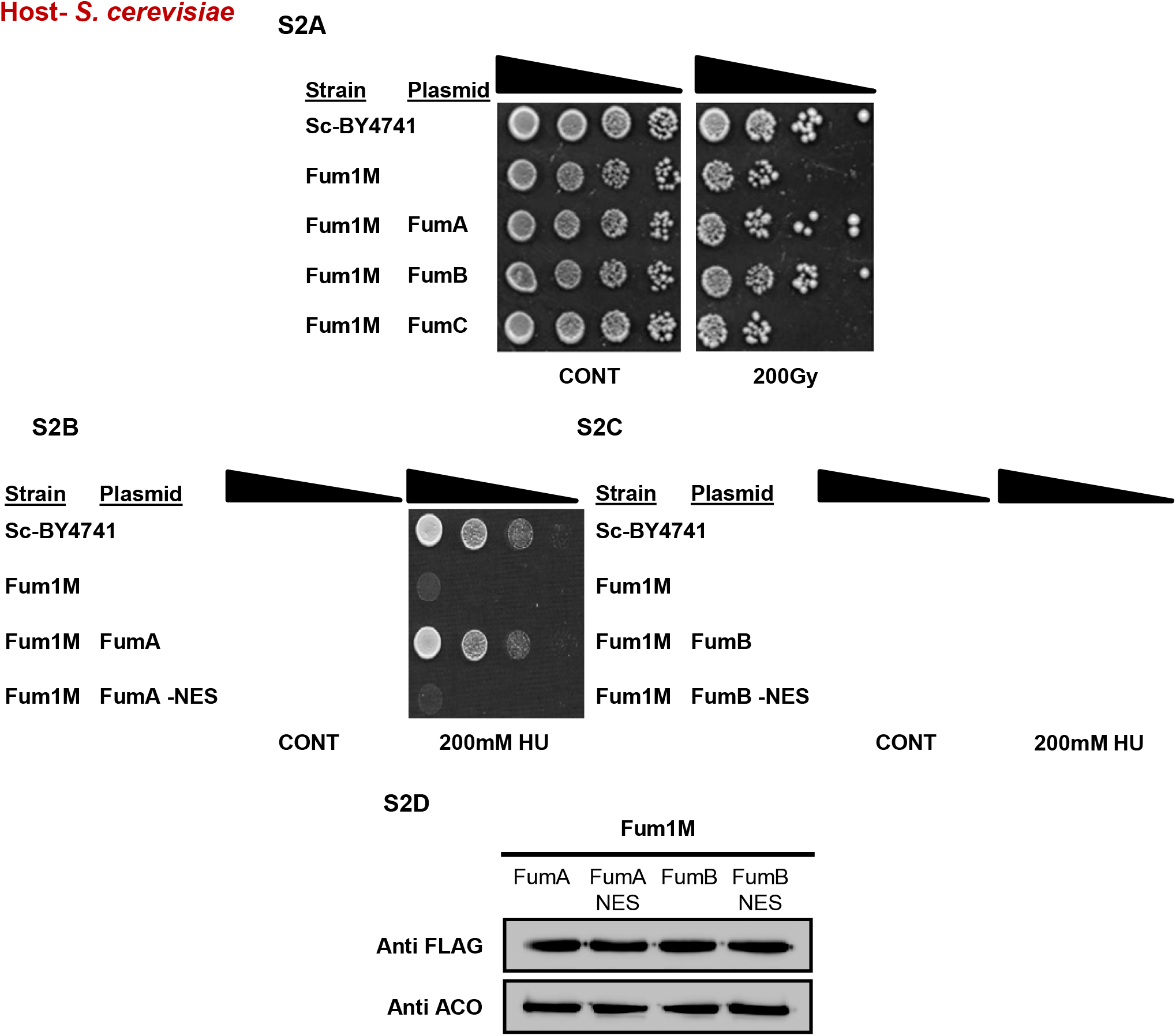
FumA-NES and FumB-NES do not complement DNA damage sensitivity in *S. cerevisiae*. (**A, B, C**) *S. cerevisiae* wild type Sc-BY4741, *Fum1M* and *Fum1M* harboring plasmids encoding the indicated *E. coli fum* genes, were grown to exponential phase in SC-Gal medium and irradiated or serially diluted onto SC-Dex or SC-Dex+ 200 mM HU. (**C**) These strains were also lysed and centrifuged to obtain the supernatant and extracts were subjected to Western blotting, using the indicated antibodies (anti Aco1 as the loading control). Each result is representative of at least three independent experiments.

**Figure S3.**
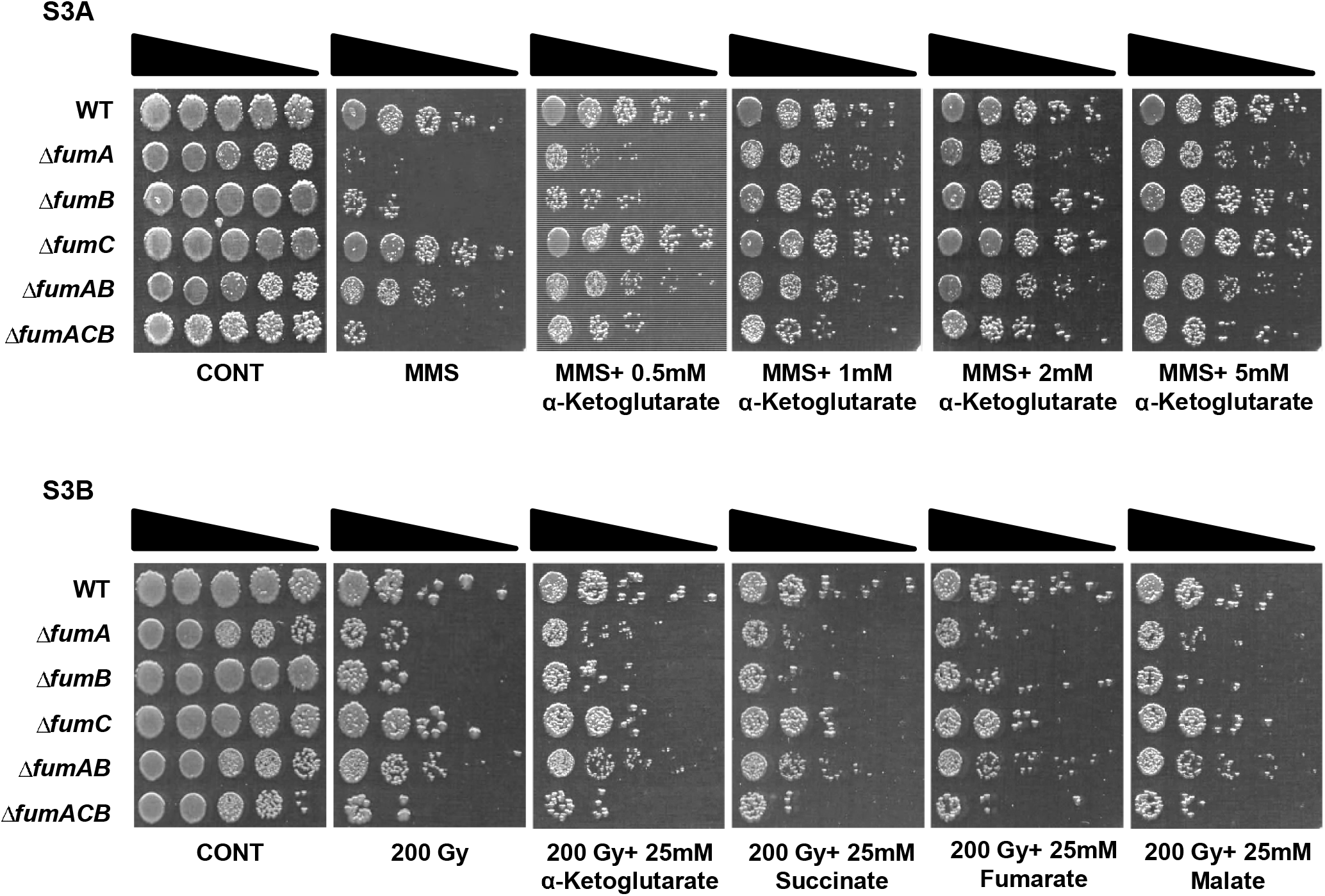
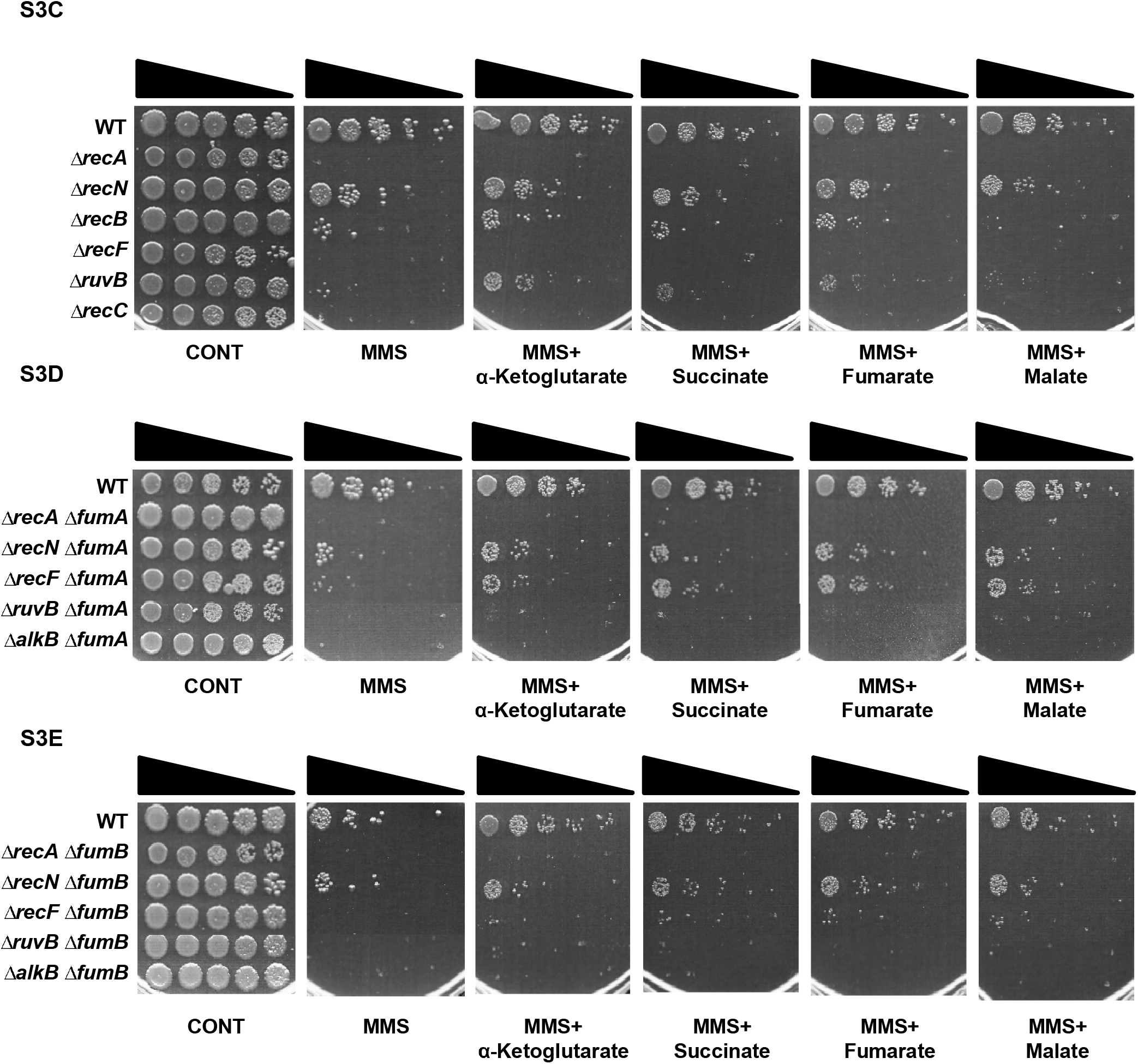
α-KG complementation of DNA damage sensitivity is specific to *fum* null mutants following MMS treatment. *E. coli* wild type (WT), *ΔfumA, ΔfumB, ΔfumC, ΔfumAB* and *ΔfumACB* strains were grown to mid-exponential phase (OD600nm=0.3) and (**A**) treated with MMS (0.35% MMS (v/v) for 45 min) <u>or</u> (**B**) the cells were irradiated (200Gy). The cells were then washed and serially diluted (spot test) onto LB plates containing or lacking the indicated organic acids. (**C**) *E. coli* DNA damage repair genes null mutants, and (**D**) composite null mutants, *ΔfumA+* DNA damage repair genes, or (**E**) *ΔfumB+* DNA damage repair genes mutants, were grown to early-exponential phase (OD600nm=0.3), and treated with MMS (0.35% (v/v) for 45 minutes at 37□C). The cells were then washed and serially diluted (spot test) onto LB plates and LB containing the indicated organic acid plates. Each result is representative of three independent experiments.

**Figure S4.**
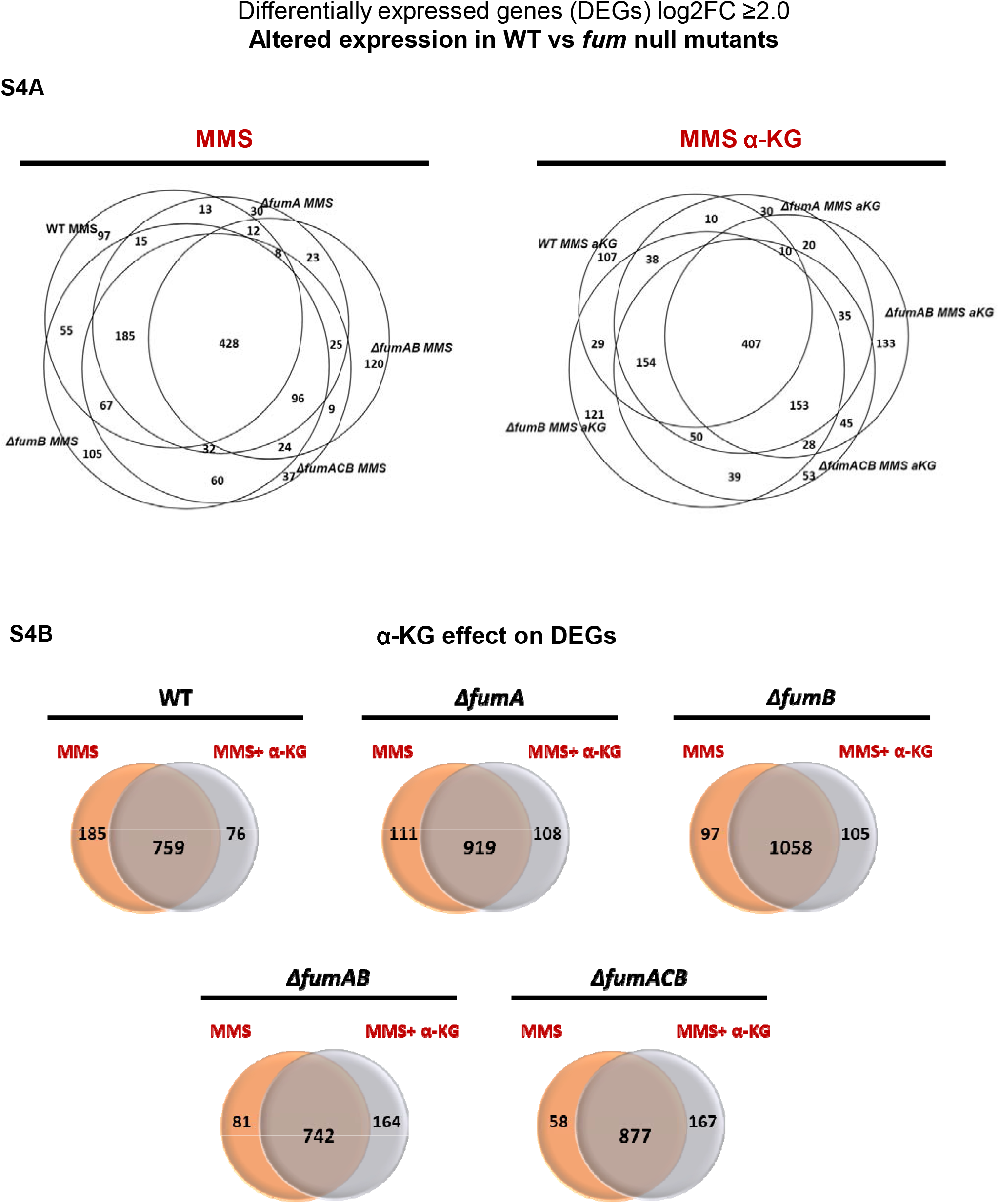
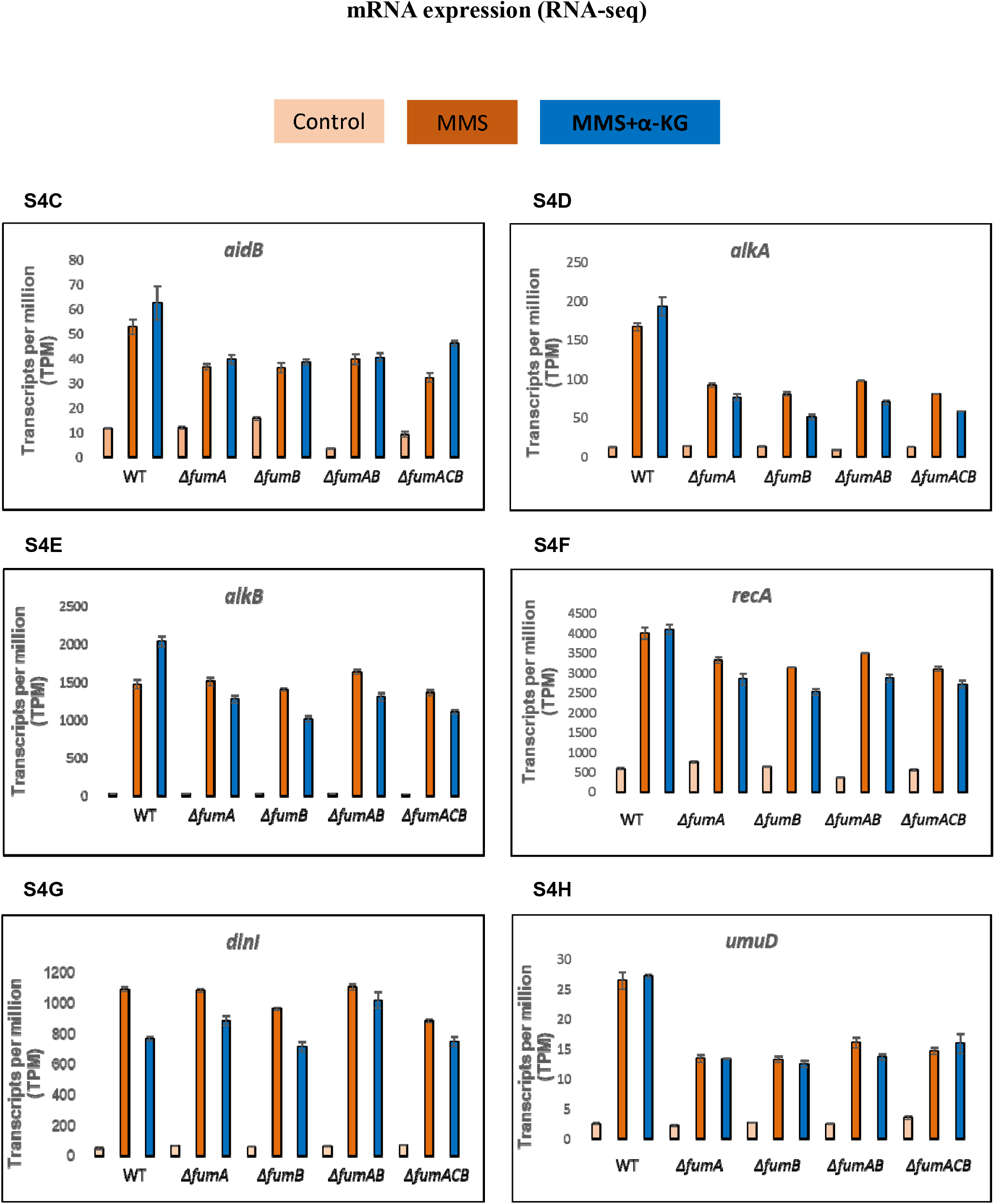
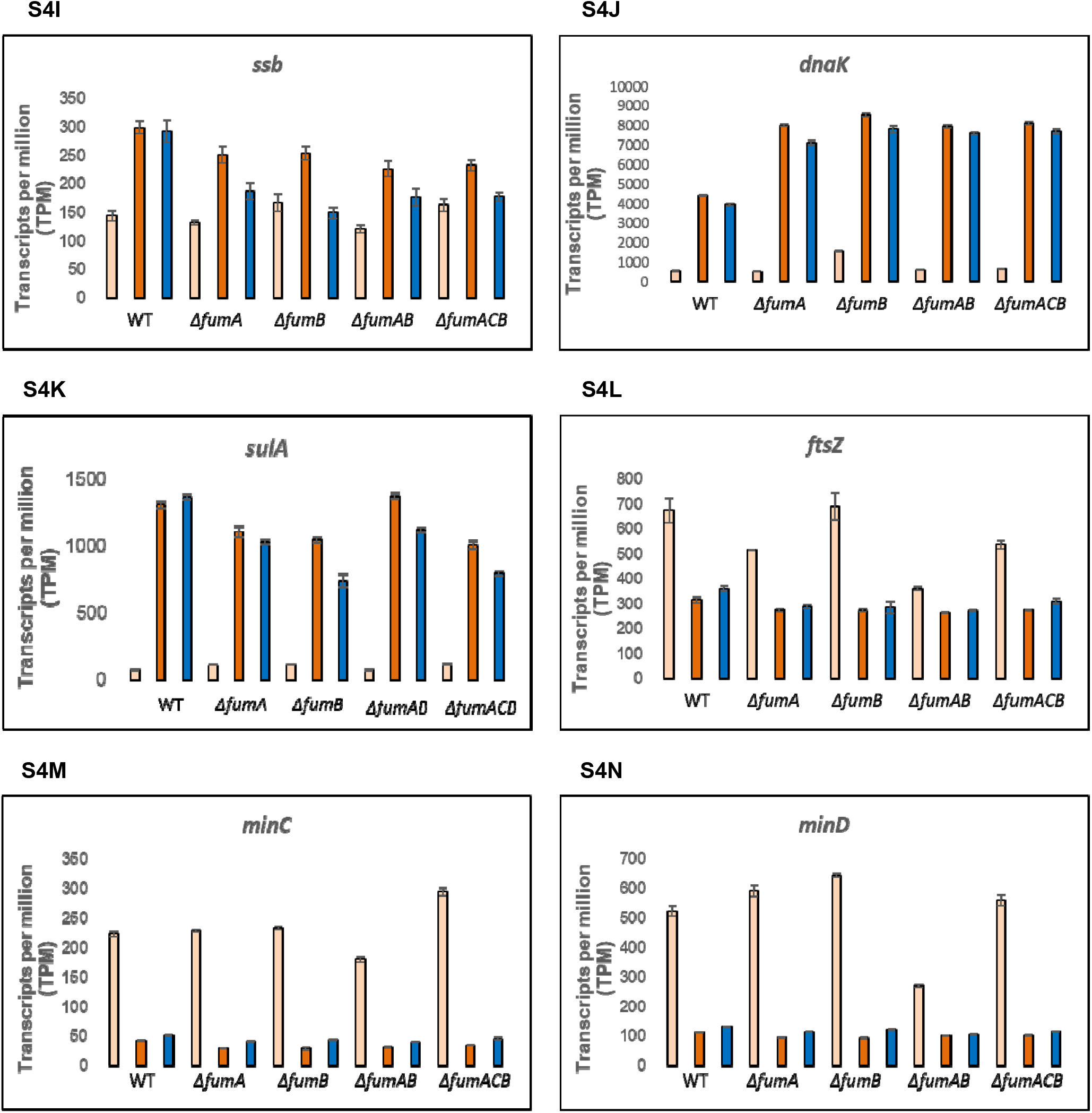
RNA-seq analysis of differential expressed genes. RNA-seq analysis; *E. coli* wild type (WT), *ΔfumA, ΔfumB, ΔfumAB* and *ΔfumACB* strains were grown to early-exponential phase (OD600nm=0.3), and treated with MMS (0.35% MMS (v/v) for 30 min). The cells were collected and total RNA was extracted and subjected to RNA-seq analysis (see materials and methods). (**A**) Venn diagrams showing the common and differential genes (transcripts) between *E. coli* wild type (WT), *ΔfumA, ΔfumB, ΔfumAB* and *ΔfumACB* strains following treatment with MMS or MMS+25mM α-KG. (**B**) comparison of shared transcripts in the presence or absence of α-KG *in E. coli* wild type (WT), *ΔfumA, ΔfumB, ΔfumAB* and *ΔfumACB* strains. (**C-N**) Gene expression levels revealed by RNA-seq for (**C**) *aidB*, Putative acyl-CoA dehydrogenase AidB (**D**) *alkA*, DNA-3-methyladenine glycosylase 2 (**E**) *alkB*, Alpha-ketoglutarate-dependent dioxygenase AlkB (**F**) *recA*, Protein RecA (**G**) *dinI*, DNA damage-inducible protein I (**H**) *umuD*, Protein UmuD (**I**) *ssb*, singl-estranded DNA-binding protein (**J**) *dnaK*, Chaperone protein DnaK (**K**) *sulA*, Cell division inhibitor SulA (**L**) *ftsZ*, Cell division protein FtsZ (**M**) *minC*, Septum site-determining protein MinC (**N**) *minD*, Septum site-determining protein *MinD*. TPM (transcripts per kilobase million) shown as Normalized gene expression from RNA-seq are mean ± standard error (n=3).

**Figure S5.**
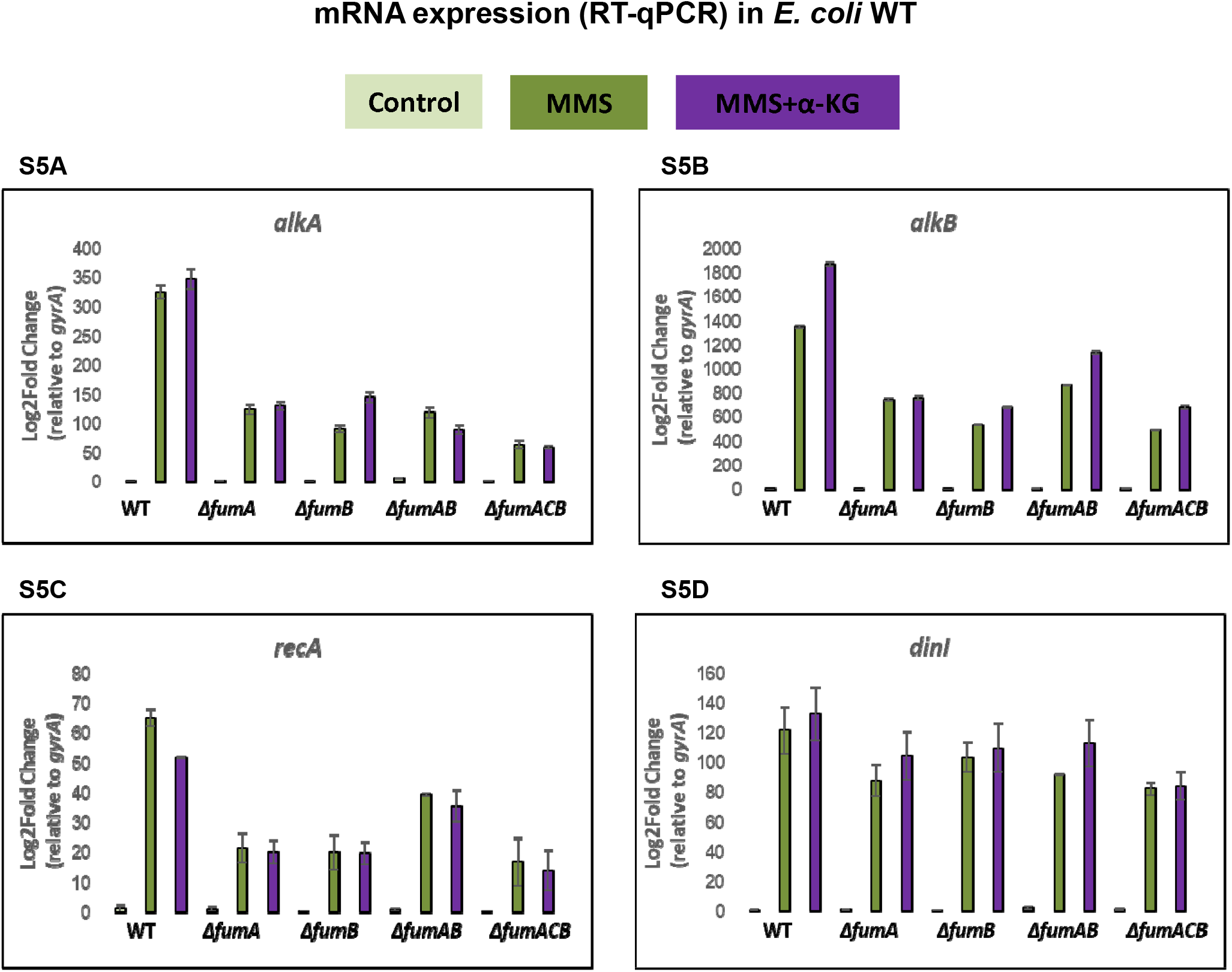
Validation of RNA-seq data. (**A-D**) *E. coli* wild type (WT), *ΔfumA, ΔfumB, ΔfumAB* and *ΔfumACB* strains were grown in LB and LB+25mM α-KG to early-exponential (OD600nm=0.3) phase, (OD 600nm=0.3), and treated with 0.35% [v/v] MMS for 35 minutes at 37□C. Cells were collected and total RNA was extracted. Graphs show the transcript levels of (**A**) *alkA* (**B**) *alkB* (**C**) *recA* and (**D**) *dinI* normalized against *gyrA* as an internal control and calculated using the ΔΔCT method. Data is represented as mean ± standard error (*n=3*).

**Figure S6.**
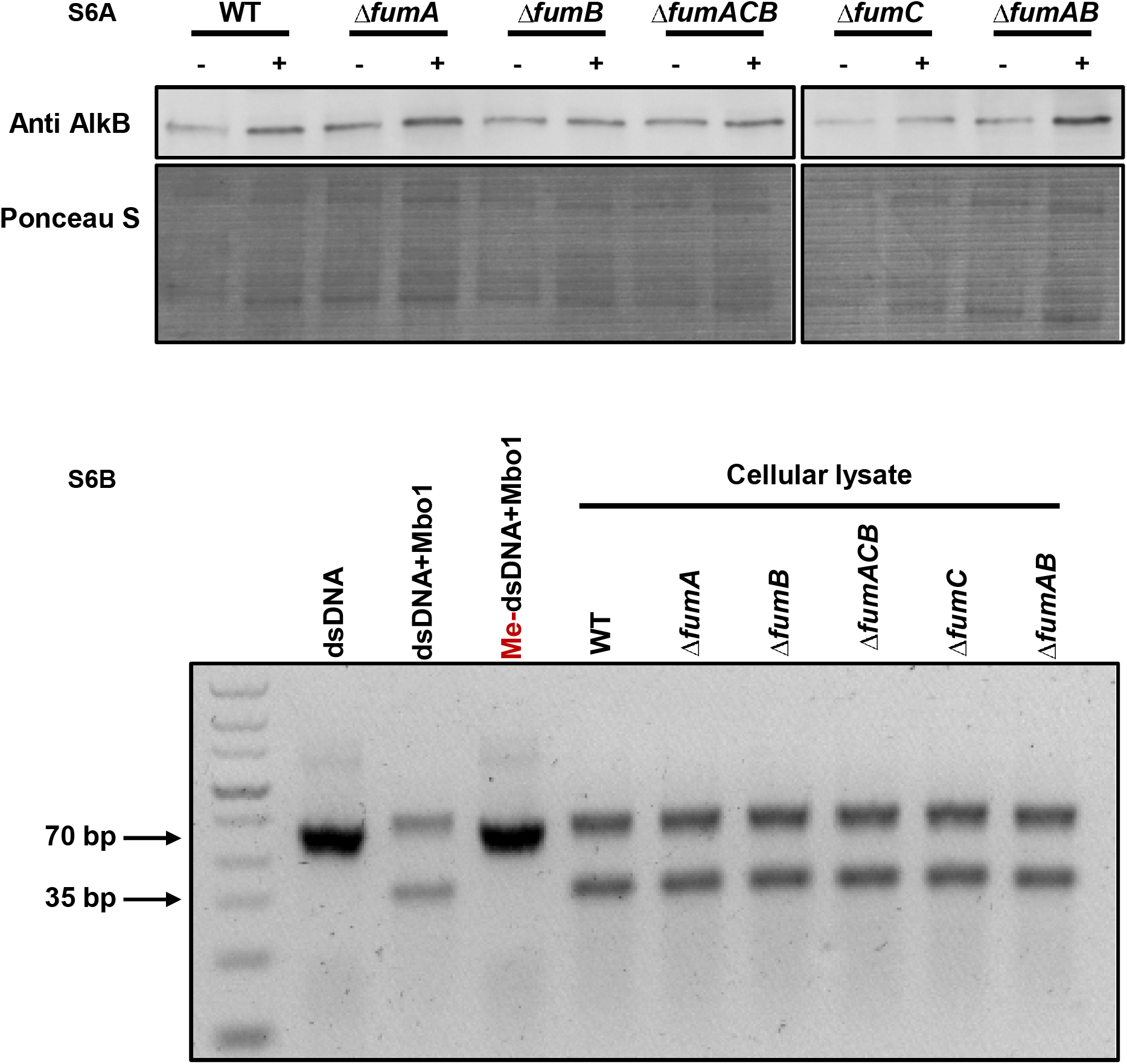
AlkB activity in *fum* null mutant strains. **(A)** *E. coli* wild type (WT), *ΔfumA, ΔfumB, ΔfumC, ΔfumAB* and *ΔfumACB* strains were grown to midexponential phase (OD600nm=0.3) and treated with MMS (0.35% MMS (v/v) for 45 min). The cells were lysed and centrifuged to obtain a supernatant which was subjected AlkB in vitro repair assay, Western blotting (using the indicated antibodies) and Ponceau S staining. **(B)** Repair of methylated oligonucleotides for 4 hours at 37□C by 40μM lysate from the indicated strains. Following in vitro repair, the reaction was mixed with equi-molar concentration of complementary ssDNA, the resultant dsDNA was digested with MboI and subjected to agarose electrophoresis and staining with safeU. Following successful demethylation, digestion with MboI produces a 35bp product.

## References

1. Woods, S.A., S.D. Schwartzbach, and J.R. Guest, Two biochemically distinct classes of fumarase in Escherichia coli. Biochim Biophys Acta, 1988. 954(1):p. 14–26.

2. Akiba, T., K. Hiraga, and S. Tuboi, Intracellular distribution of fumarase in various animals. J Biochem, 1984. 96(1):p. 189–95.

3. Tuboi, S., et al., Rat liver mitochondrial and cytosolic fumarases with identical amino acid sequences are encoded from a single mRNA with two alternative in-phase AUG initiation sites. Adv Enzyme Regul, 1990. 30:p. 289–304.

4. Dik, E., et al., Human Fumarate Hydratase Is Dual Localized by an Alternative Transcription Initiation Mechanism. Traffic, 2016. 17(7): p. 720–32.

5. Karniely, S., N. Regev-Rudzki, and O. Pines, The presequence of fumarase is exposed to the cytosol during import into mitochondria. J Mol Biol, 2006. 358(2):p. 396–405.

6. Pracharoenwattana, I., et al., Arabidopsis has a cytosolic fumarase required for the massive allocation of photosynthate into fumaric acid and for rapid plant growth on high nitrogen. Plant J, 2010. 62(5): p. 785–95.

7. Regev-Rudzki, N., O. Yogev, and O. Pines, The mitochondrial targeting sequence tilts the balance between mitochondrial and cytosolic dual localization. J Cell Sci, 2008. 121(Pt 14): p. 2423–31.

8. Sass, E., S. Karniely, and O. Pines, Folding of fumarase during mitochondrial import determines its dual targeting in yeast. J Biol Chem, 2003. 278(46): p. 45109–16.

9. Yogev, O., et al., Fumarase: a mitochondrial metabolic enzyme and a cytosolic/nuclear component of the DNA damage response. PLoS Biol, 2010. 8(3): p. e1000328.

10. Jiang, Y., et al., Author Correction: Local generation of fumarate promotes DNA repair through inhibition of histone H3 demethylation. Nat Cell Biol, 2018. 20(10): p. 1226.

11. Leshets, M., et al., Fumarase is involved in DNA double-strand break resection through a functional interaction with Sae2. Curr Genet, 2018. 64(3):p. 697–712.

12. Singer, E., et al., Bacterial fumarase and L-malic acid are evolutionary ancient components of the DNA damage response. Elife, 2017. 6.

13. van Vugt-Lussenburg, B.M., et al., Biochemical similarities and differences between the catalytic [4Fe-4S] cluster containing fumarases FumA and FumB from Escherichia coli. PLoS One, 2013. 8(2): p. e55549.

14. Tseng, C.P., Regulation of fumarase (fumB) gene expression in Escherichia coli in response to oxygen, iron and heme availability: role of the arcA, fur, and hemA gene products. FEMS Microbiol Lett, 1997. 157(1):p. 67–72.

15. Park, S.J. and R.P. Gunsalus, Oxygen, iron, carbon, and superoxide control of the fumarase fumA and fumC genes of Escherichia coli: role of the arcA, fnr, and soxR gene products. J Bacteriol, 1995. 177(21): p. 6255–62.

16. Burak, E., et al., Evolving dual targeting of a prokaryotic protein in yeast. Mol Biol Evol, 2013. 30(7):p. 1563–73.

17. Feliciano, P.R., C.L. Drennan, and M.C. Nonato, Crystal structure of an Fe-S cluster-containing fumarate hydratase enzyme from Leishmania major reveals a unique protein fold. Proc Natl Acad Sci U S A, 2016. 113(35): p. 9804–9.

18. Baharoglu, Z. and D. Mazel, SOS, the formidable strategy of bacteria against aggressions. FEMS Microbiol Rev, 2014. 38(6): p. 1126–45.

19. Errol, C.F., et al., DNA Repair and Mutagenesis, Second Edition. 2006: American Society of Microbiology.

20. Kreuzer, K.N., DNA damage responses in prokaryotes: regulating gene expression, modulating growth patterns, and manipulating replication forks. Cold Spring Harb Perspect Biol, 2013. 5(11): p. a012674.

21. The Gene Ontology Resource: 20 years and still GOing strong. Nucleic Acids Res, 2019. 47(D1): p. D330–d338.

22. Ashburner, M., et al., Gene ontology: tool for the unification of biology. The Gene Ontology Consortium. Nat Genet, 2000. 25(1):p. 25–9.

23. Kronenberg, J., et al., Fumaric Acids Directly Influence Gene Expression of Neuroprotective Factors in Rodent Microglia. Int J Mol Sci, 2019. 20(2).

24. Sciacovelli, M. and C. Frezza, Fumarate drives EMT in renal cancer. Cell Death Differ, 2017. 24(1):p. 1–2.

25. Sciacovelli, M., et al., Corrigendum: Fumarate is an epigenetic modifier that elicits epithelial-to-mesenchymal transition. Nature, 2016. 540(7631): p. 150.

26. Falnes, P.O., A. Klungland, and I. Alseth, Repair of methyl lesions in DNA and RNA by oxidative demethylation. Neuroscience, 2007. 145(4): p. 1222–32.

27. Fedeles, B.I., et al., The AlkB Family of of Fe(II)/α-Ketoglutarate-dependent Dioxygenases: Repairing Nucleic Acid Alkylation Damage and Beyond. J Biol Chem, 2015. 290(34): p. 20734–42.

28. Xiao, M., et al., Inhibition of alpha-KG-dependent histone and DNA demethylases by fumarate and succinate that are accumulated in mutations of FH and SDH tumor suppressors. Genes Dev, 2012. 26(12):p. 1326–38.

29. Shivange, G., N. Kodipelli, and R. Anindya, A nonradioactive restriction enzyme-mediated assay to detect DNA repair by Fe(II)/2-oxoglutarate-dependent dioxygenase. Anal Biochem, 2014. 465:p. 35–7.

30. Shivange, G., et al., RecA stimulates AlkB-mediated direct repair of DNA adducts. Nucleic Acids Res, 2016. 44(18): p. 8754–8763.

31. Johnson, T.I., et al., Fumarate hydratase loss promotes mitotic entry in the presence of DNA damage after ionising radiation. Cell Death Dis, 2018. 9(9): p. 913.

32. Hausinger, R.P., FeII/alpha-ketoglutarate-dependent hydroxylases and related enzymes. Crit Rev Biochem Mol Biol, 2004. 39(1):p. 21–68.

33. Loenarz, C. and C.J. Schofield, Expanding chemical biology of 2-oxoglutarate oxygenases. Nat Chem Biol, 2008. 4(3): p. 152–6.

34. Gottlieb, E. and I.P. Tomlinson, Mitochondrial tumour suppressors: a genetic and biochemical update. Nat Rev Cancer, 2005. 5(11): p. 857–66.

35. Pollard, P.J., et al., Accumulation of Krebs cycle intermediates and over-expression of HIF1alpha in tumours which result from germline FH and SDH mutations. Hum Mol Genet, 2005. 14(15): p. 2231–9.

36. Selak, M.A., et al., Succinate links TCA cycle dysfunction to oncogenesis by inhibiting HIF-alpha prolyl hydroxylase. Cancer Cell, 2005. 7(1): p. 77–85.

37. Tahiliani, M., et al., Conversion of 5-methylcytosine to 5-hydroxymethylcytosine in mammalian DNA by MLL partner TET1. Science, 2009. 324(5929): p. 930–5.

38. Keyamura, K., et al., RecA protein recruits structural maintenance of chromosomes (SMC)-like RecN protein to DNA double-strand breaks. J Biol Chem, 2013. 288(41):p. 29229–37.

39. Datsenko, K.A. and B.L. Wanner, One-step inactivation of chromosomal genes in Escherichia coli K-12 using PCR products. Proc Natl Acad Sci U S A, 2000. 97(12): p. 6640–5.

40. Ben-Menachem, R., et al., Yeast aconitase mitochondrial import is modulated by interactions of its C and N terminal domains and Ssa1/2 (Hsp70). Sci Rep, 2018. 8(1): p. 5903.

41. Yang, J., et al., The I-TASSER Suite: protein structure and function prediction. Nat Methods, 2015. 12(1): p. 7–8.

42. Bohnuud, T., et al., Detection of Peptide-Binding Sites on Protein Surfaces Using the Peptimap Server. Methods Mol Biol, 2017. 1561:p. 11–20.

43. Ashkenazy, H., et al., ConSurf 2016: an improved methodology to estimate and visualize evolutionary conservation in macromolecules. Nucleic Acids Res, 2016. 44(W1):p. W344–50.

44. Chen, S., et al., fastp: an ultra-fast all-in-one FASTQ preprocessor. Bioinformatics, 2018. 34(17): p. i884–i890.

45. Li, H. and R. Durbin, Fast and accurate short read alignment with Burrows-Wheeler transform. Bioinformatics, 2009. 25(14): p. 1754–60.

46. Pertea, M., et al., StringTie enables improved reconstruction of a transcriptome from RNA-seq reads. Nat Biotechnol, 2015. 33(3): p. 290–5.

47. Kruger, N.J., The Bradford method for protein quantitation. Methods Mol Biol, 1994. 32:p. 9–15.

48. Bradford, M.M., A rapid and sensitive method for the quantitation of microgram quantities of protein utilizing the principle of protein-dye binding. Analytical biochemistry, 1976. 72:p. 248–254.

