## Supplemental tables for "A novel combination of Class-I fumarases and metabolites (α-ketoglutarate and fumarate) signal the DNA damage response in *E. coli*" (from another version of this preprint: version 4, 2020-12-27)

**Supplementary table 1- List of differentially expressed genes expressed in either the WT strain, or in the *fum* null mutant strains.**

| **Exclusive to WT MMS** | **Common elements in *ΔfumA* and *ΔfumB* MMS (not in the other strains)** | **Exclusive to WT MMS α-KG** | **Common elements in *ΔfumA* and *ΔfumB* MMS α-KG (not in the other strains)** | **Exclusive to *ΔfumAB* MMS** | **Exclusive to *ΔfumAB* MMS α-KG** | **Exclusive to *ΔfumACB* MMS** | **Exclusive to *ΔfumACB* MMS α-KG** |
|---|---|---|---|---|---|---|---|
| aceF | aceA | aceB | actP | ahpC | ahpC | dosC | atoD |
| ackA | aceK | aceE | agrA | arsC | argO | eutN | casC |
| ade | acs | aceF | alsB | cfa | arsC | frvA | cusB |
| adiC | actP | ackA | amyA | citF | blc | glcE | dosC |
| agaV | ais | ade | argU | clpA | bolA | ileX | elaD |
| araC | alaE | adiC | chbB | copA | citF | kdpB | eutE |
| aroM | argS | agaV | citT | cpsB | clpA | mngA | eutN |
| betT | argU | araC | csiD | cueO | cpsB | php | fepE |
| bfr | bdm | araH | dadA | dcyD | cysQ | ydiO | frvA |
| bglF | bglG | argH | dadX | eutS | dsrA | ygbA | glcE |
| cadB | csgD | argT | degP | fixA | ecnB | yhdV | ileX |
| can | cusF | aroM | dppB | fliY | emrY | yihP | mngA |
| citE | dadA | arrS | dppC | fsaB | fixA | yihQ | ompG |
| curA | dadX | betA | dppF | hchA | fixC | | php |
| dsdC | degP | betT | ecnA | hiuH | fliY | | uidC |
| dsrA | dppB | bfr | fadB | hlyE | gloA | | wcaI |
| exbB | dppC | can | fdhE | hspQ | hchA | | ybbY |
| exbD | dppF | cdh | fdoG | ilvC | hiuH | | yceJ |
| fbaA | fadB | citE | fimB | ldtE | hlyE | | ydiO |
| fhlA | fdhE | cysM | flgK | osmC | hspQ | | ydjJ |
| frwA | fdoG | dhaK | gabD | paaZ | htpX | | yfbO |
| fumB | fimB | dsdC | gabP | poxB | ilvC | | yhdV |
| gadB | gabD | epmA | gabT | rutF | ldtE | | yihP |
| gadE | gabP | evgS | glgP | sokC | osmC | | yihQ |
| gapA | glgX | exbB | gltA | torY | poxB | | |
| glsA | gltA | exbD | glyY | yafN | psiF | | |
| gudD | glyU | frwA | gspA | yaiT | pspA | | |
| holE | glyX | fsaA | gspG | ycgY | pyrB | | |
| ibaG | gntU | fumB | hupA | ydhR | pyrI | | |
| idnK | hofC | gadE | intF | yedK | rclR | | |
| inaA | hupA | glsA | leuQ | yedL | rutF | | |
| ldtC | intF | gltB | lhgO | yedP | torY | | |
| livG | lhgO | gntP | lldP | yegP | ulaG | | |
| lpxT | lldP | gudD | lsrB | yfcP | uspB | | |

| | | | | | |
|---|---|---|---|---|---|
| menH | lsrF | hcp | lsrF | yffR | yafL |
| metF | lsrG | hlyE | lsrG | yfiP | yahC |
| mgrR | paaA | hyaA | maeB | yhcO | yaiT |
| mgtA | paaB | idnK | mglA | yiaK | ybaY |
| micF | paaC | ilvG | mlaF | yibH | ybfI |
| mlc | paaD | ilvX | nagK | yibI | ycaC |
| mntP | paaE | inaA | paaA | yibT | ycgY |
| pgsA | paaH | insH1 | paaB | yidL | yciE |
| phoA | paaK | kdgR | paaC | yidP | ydbD |
| ppdD | pepN | leuB | paaD | yjcC | ydhR |
| puuP | priB | leuD | paaE | yjdK | yeaI |
| queA | putA | lpxT | paaH | yqhD | yedL |
| queD | putP | lrhA | paaK | yqjG | yedP |
| rob | pyrH | lsrR | patA | ysaD | yegP |
| rraA | recF | metF | peaD | | yfcP |
| rrfC | rhaD | micF | pepN | | yffQ |
| rrfD | rplQ | mlc | pheL | | yffR |
| rrfG | rpmD | mocA | ppsA | | yfjP |
| rsxA | rpoA | nlpA | ppsR | | yfjQ |
| sokE | rpsE | nrfE | putA | | ygaU |
| thrA | rtcR | omrB | putP | | yghA |
| topA | sdhB | phoA | recF | | yiaK |
| tqsA | sdhC | ppdD | rplF | | yiaM |
| trpL | sra | pphA | rplO | | yibH |
| uhpB | tap | pqqL | rplQ | | yibI |
| uhpC | tar | prlF | rplR | | yibT |
| ushA | tfaE | proX | rpmD | | yidL |
| uspG | tfaP | puuA | rpsE | | yjbJ |
| yagU | uidB | puuD | rpsN | | yjdJ |
| ybeQ | xdhB | puuP | sdhB | | yjdK |
| ybeU | yaiT | queA | sdhC | | ykgH |
| ybgO | ybcV | queD | selC | | ykgR |
| ybhS | ybfK | rob | sucB | | ymgE |
| yciE | ybhJ | rraA | tap | | yoaC |
| ydcI | ybjG | rsxA | tar | | yqaE |
| yddB | ybjM | slp | tfaE | | yqhD |
| ydhU | ycaP | thrA | thrV | | yqjG |
| ydhZ | ycfP | thrS | valV | | ysaD |
| ydiE | ycjQ | topA | yaaA | | |
| yeaD | ycjR | trpC | ybcV | | |
| yeaR | yfaZ | uhpB | ybfK | | |
| yegH | yfeO | uhpC | ybgA | | |
| yegP | yfhL | uspG | ybiX | | |
| yegZ | yfjD | xylF | ybjM | | |
| yeiB | yghQ | ybiA | ycaP | | |

| | | | |
|---|---|---|---|
| yeiR | yhhX | ycaM | ycjQ |
| yfbT | yhjE | yccM | ycjR |
| yfdY | yjbD | ydbJ | ydgA |
| yfeX | ymgJ | ydcI | yfjD |
| ygaM | ypjI | yddB | yjaA |
| ygcB | | ydeE | ykiA |
| yggX | | ydhU | yodC |
| ygjJ | | ydiE | ypjI |
| ygjQ | | ydiU | |
| yhdT | | ydjH | |
| yhfL | | ydjX | |
| yhfU | | yeaD | |
| yidE | | yeaR | |
| yihF | | yfbM | |
| yihM | | yfdY | |
| yjhZ | | yfeX | |
| ynfD | | yfjK | |
| ynfG | | ygjJ | |
| ynfO | | yhbW | |
| yobH | | yhfU | |
| yoeF | | yidE | |
| yoeH | | yihN | |
| yqgB | | yiiS | |
| ythA | | yjhZ | |
| | | ylcI | |
| | | yncG | |
| | | ynfE | |
| | | ynfF | |
| | | ynfG | |
| | | ynfO | |
| | | yohJ | |
| | | yqgB | |
| | | zraS | |

#### Supplementary table 2- Bacterial strains used in this study

| Strain | Reference |
|---|---|
| BW25113 | Coli Genetic Stock Center (CGSC) |
| (Δ*fumA*)-JW1604-1 | CGSC |
| (Δ*fumB*)-JW4083-1 | CGSC |
| (Δ*fumC*)-JW1603-1 | CGSC |
| (Δ*alkB*)- JW2200-1 | CGSC |
| (Δ*recA*)- JW2669-1 | CGSC |
| (Δ*recN*)- JW5416-1 | CGSC |
| (Δ*recF*)- JW3677-1 | CGSC |
| (Δ*recB*)- JW2788-1 | CGSC |
| (Δ*recC*)- JW2790-1 | CGSC |
| (Δ*ruvB*)- JW1849-2 | CGSC |
| Δ*fumAB* | This study |
| Δ*fumAC* | This study |
| Δ*alkB* Δ*fumA* | This study |
| Δ*alkB* Δ*fumB* | This study |
| Δ*recA* Δ*fumA* | This study |
| Δ*recA* Δ*fumB* | This study |
| Δ*recN* Δ*fumA* | This study |
| Δ*recN* Δ*fumB* | This study |
| Δ*recF* Δ*fumA* | This study |
| Δ*recF* Δ*fumB* | This study |
| Δ*ruvB* Δ*fumA* | This study |

**Supplementary table 3- Plasmids and primers used in this study**

| Name | Forward primer-5' | Reverse primer-5' |
|---|---|---|
| FumA_pGAL425 | GGATCCATGTCAAACAAACCCTTTCA | GTCGACTTA**TTTATCGTCATCGTCTTTGTAGT C**TTTCACACAGCGGGTGC |
| FumA-NES_pGAL425 | GGATCCATGCTAGAAGCGAATTGGCTCTCA AGTTGGCTGGTCTTGATATTAACG | GGATCC-TTATTTCACACAGCGGGTG |
| FumB_pGAL425 | GGATCCATGTCAAACAAACCCTTTATC | GTCGACTTA**TTTATCGTCATCGTCTTTGTAGT C** CTTAGTGCAGTTCGCGCA |
| FumB-NES_pGAL425 | GGATCCATGCTAGAAGCGAATTGGCTCTCA AGTTGGCTGGTCTTGATATTAACG | GGATCCTTACTTAGTGCAGTTCGCG |
| FumC_pGAL425 | GGATCCATGAATACAGTACGCAGCG | GTCGACTTA**TTTATCGTCATCGTCTTTGTAGT C** ACGCCCGGCTTTCATACT |
| FumC-NES_pGAL425 | GGATCCATGCTAGAAGCGAATTGGCTCTCA AGTTGGCTGGTCTTGATATTAACG | GTCGACTTA**TTTATCGTCATCGTCTTTGTAGT C** ACGCCCGGCTTTCATACT |
| FumC-NLS_pGAL425 | GGATCCATGCTAGTAGCCCTCCAAAGAAGA AGAGGAAGCTGGGTTCTAGAGCTAGCG | GTCGACTTA**TTTATCGTCATCGTCTTTGTAGT C** ACGCCCGGCTTTCATACT |
| FumC-mNLS_pGAL425 | GGATCCATGCTAAGCGCCCTCCAAGAGGA TGAGGGAAGCTGGGTTCTAAGACTAGCG | GTCGACTTA**TTTATCGTCATCGTCTTTGTAGT C** ACGCCCGGCTTTCATACT |
| FumA_PUC18 | GGATCCATGTCAAACAAACCCTTTATC | GTCGACTTA**TTTATCGTCATCGTCTTTGTAGT C** CTTAGTGCAGTTCGCGCA |
| FumA-Exc (Y481D) PUC18 | GACCTCGGTAGCATCGGC | GAAGCCGCCGTGTTTATG |
| FumA-Exc (1233K T236I) PUC18 | AAGGCGGTATTTCTGCGGAAACCAACCTG | TCACAAACGCGATATGGAAC |
| FumB_PUC18 | GGATCCATGCTAGAAGCGAATTGGCTCTCA AGTTGGCTGGTCGATATTAACG | GGATCCTTACTTAGTGCAGTTCGCG |
| FumB-Exc (Y481D) PUC18 | GGATCCATGAATACAGTACGCAGCG | GTCGACTTA**TTTATCGTCATCGTCTTTGTAGT C**ACGCCCGGCTTTCATACT |
| FumB-Exc (1233K T236I) PUC18 | GGATCCATGCTAGAAGCGAATTGGCTCTCA AGTTGGCTGGTCTTGATATTAACG | GTCGACTTA**TTTATCGTCATCGTCTTTGTAGT C** ACGCCCGGCTTTCATACT |
| PKD46, PKD3, PKD4 | Kind gift from Ilan Rosenshine | |

### for all FLAG tagged constructs, the FLAG sequence **TTTATCGTCATCGTCTTTGTAGTC** was introduced within the primer.

#### Supplementary table 4- Yeast strains used in this study

| Strain | Reference |
|---|---|
| BY4741 | Personal stock |
| *Δfum1/cox2::*FUM1$^{m}$ (referred to as Fum1M) | Yogev O. 2010 |
| Δ*fum1* | Sass E et al. 2003 |

#### Supplementary table 5- RNA-seq quality control

| Sample_id | Mapped_reads | Mapped_ratio |
|---|---:|---:|
| BW251152-WT1 | 13026168 | 99.90% |
| BW251152-WT2 | 25631302 | 97.40% |
| BW251152-WT3 | 65172557 | 97.70% |
| BW251153-MMS1 | 10332122 | 100% |
| BW251153-MMS2 | 17609977 | 99.90% |
| BW251153-MMS3 | 29223633 | 96.10% |
| BW251153-MMSAKG-1 | 14836645 | 99.90% |
| BW251153-MMSAKG-2 | 25725731 | 97.10% |
| BW251153-MMSAKG-3 | 25664599 | 96.80% |
| ΔfumA-1 | 15822922 | 99.80% |
| ΔfumA-2 | 10943808 | 99.80% |
| ΔfumA-3 | 33930739 | 95.10% |
| ΔfumA-MMS1 | 10630673 | 99.80% |
| ΔfumA-MMS2 | 14093714 | 99.80% |
| ΔfumA-MMS3 | 22169461 | 98.60% |
| ΔfumA-MMSAKG-1 | 10426749 | 99.60% |
| ΔfumA-MMSAKG-2 | 22506224 | 94.60% |
| ΔfumA-MMSAKG-3 | 48982551 | 94.40% |
| ΔfumAB-1 | 10460591 | 99.60% |
| ΔfumAB-2 | 24015721 | 99.40% |
| ΔfumAB-3 | 21882329 | 98% |
| ΔfumAB-MMS1 | 9509178 | 99.70% |
| ΔfumAB-MMS2 | 17954479 | 99.30% |
| ΔfumAB-MMS3 | 17356595 | 97.30% |
| ΔfumAB-MMSAKG-1 | 11107770 | 99.40% |
| ΔfumAB-MMSAKG-2 | 25421689 | 94.40% |
| ΔfumAB-MMSAKG-3 | 91308557 | 98.70% |
| ΔfumABC-1 | 9760685 | 99.70% |
| ΔfumABC-2 | 25214587 | 99.60% |

| | | |
|---|---|---|
| ΔfumABC-3 | 35968295 | 95.40% |
| ΔfumABC-MMS1 | 12038975 | 99.70% |
| ΔfumABC-MMS2 | 22783437 | 95% |
| ΔfumABC-MMS3 | 34782735 | 96.20% |
| ΔfumABC-MMSAKG-1 | 10879818 | 99.30% |
| ΔfumABC-MMSAKG-2 | 14295702 | 96.90% |
| ΔfumABC-MMSAKG-3 | 26650790 | 98.10% |
| ΔfumB-1 | 13616787 | 99.80% |
| ΔfumB-2 | 16714019 | 99.80% |
| ΔfumB-3 | 39885293 | 97.80% |
| ΔfumB-MMS1 | 5810281 | 99.70% |
| ΔfumB-MMS2 | 42810691 | 99.60% |
| ΔfumB-MMS3 | 20484749 | 97.40% |
| ΔfumB-MMSAKG-1 | 14252956 | 99.50% |
| ΔfumB-MMSAKG-2 | 26020671 | 95.20% |
| ΔfumB-MMSAKG-3 | 50677026 | 98.80% |
